# Investigation of heterochromatin protein 1 function in the malaria parasite *Plasmodium falciparum* using a conditional domain deletion and swapping approach

**DOI:** 10.1101/2020.11.26.400325

**Authors:** Hai T. N. Bui, Armin Passecker, Nicolas Brancucci, Till S. Voss

## Abstract

The human malaria parasite *Plasmodium falciparum* encodes a single ortholog of heterochromatin protein 1 (PfHP1) that plays a crucial role in the epigenetic regulation of various survival-related processes. PfHP1 is essential for parasite proliferation and the heritable silencing of genes linked to antigenic variation, host cell invasion and sexual conversion. Here, we employed CRISPR/Cas9-mediated genome editing combined with the DiCre/LoxP system to investigate how the PfHP1 chromodomain (CD), hinge domain and chromoshadow domain (CSD) contribute to overall PfHP1 function. We show that the C-terminal 76 residues are responsible for targeting PfHP1 to the nucleus. Furthermore, we reveal that each of the three functional domains of PfHP1 are required for heterochromatin formation, gene silencing and mitotic parasite proliferation. Finally, we discovered that the hinge and CSD domains of HP1 are functionally conserved between *P. falciparum* and *P. berghei*, a related malaria parasite infecting rodents. In summary, our study provides new insights into PfHP1 function and offers a tool for further studies on epigenetic regulation and life cycle decision in malaria parasites.

## Introduction

Heterochromatin protein 1 (HP1) was initially described in *Drosophila melanogaster* as a non-histone chromosomal protein associated with heterochromatin and responsible for variegated gene expression (1, 2). By binding to the repressive tri-methylation mark on lysine 9 of histone 3 (H3K9me3) (3, 4) HP1 facilitates the formation and spreading of heterochromatin. Chromatin-bound HP1 serves as a platform for the recruitment of downstream chromatin modifiers including H3K9-specific histone SU(VAR)3-9-like lysine methyltransferases (HKMT) that methylate H3K9 on neighboring nucleosomes and thus facilitate the binding of further HP1 proteins (5, 6). The self-propagation of HP1 results in the regional spreading of heterochromatin, thereby promoting silencing of heterochromatin-associated genes (7–10). HP1 is a small protein and well conserved among eukaryotes (11). Orthologs have been identified in a broad range of unicellular and multicellular organisms (7, 11) including the evolutionary divergent protozoan parasites of the genus *Plasmodium*, the causative agents of malaria (12, 13). *S. pombe* contains two members of the HP1 protein family, namely Switching 6 (Swi6) and Chromodomain-containing protein 2 (Chp2), three variants (HP1 α, HP1 β, HP1γ) are encoded in the mammalian genome and D. melanogaster possesses five HP1 variants (HP1a-e) (8, 11). HP1 proteins consist of three distinct functional domains: a conserved chromodomain (CD) at the N-terminus, which binds H3K9me3 (3, 4); a conserved chromoshadow domain (CSD) at the C-terminus, which mediates HP1 dimerization and interaction with other chromatin modifiers (5, 14–17); and a variable hinge region separating the CD and CSD and shown to interact with histone H1, DNA or RNA (18–21).

In mammals, chromatin localization of HP1 proteins has been shown to be variant-specific. While mammalian HP1α and HP1β are primarily found in pericentromeric heterochromatin, HP1γ is found in both heterochromatic and euchromatic regions (22–25). The requirements for targeting HP1 to heterochromatin show inter-species differences. In fission yeast, the CD domain was shown to direct Swi6 to heterochromatin (26), whereas in mice this function additionally requires RNA binding by the hinge domain (19). In *D. melanogaster* HP1 the N-terminal 95 residues encompassing the CD and the C-terminal residues 95-206 containing the hinge domain and CSD both target HP1 to pericentromeric heterochromatin (27, 28). Similarly, HP1 domains play different roles in targeting HP1 to the nucleus in different species. In *D. melanogaster*, the C-terminal 54 residues of the protein (amino acids 152-206) are required for importing HP1 to the nucleus (27). In *S. pombe*, however, the hinge region of Swi6 plays a dominant role in directing Swi6 to the nucleus. Additionally, the Swi6 C-terminus acts as a second, albeit weaker, nuclear targeting domain (26).

Malaria parasites possess only a single HP1 ortholog that primarily binds to chromosomal regions containing members of multigene families encoding variant surface antigens (12, 13, 29, 30). Unlike its orthologs in other eukaryotes (31–34), however, HP1 is absent from pericentromeric regions in malaria parasites and plays no apparent role in the formation and maintenance of centromere structure and function (12, 29, 30). *In P. falciparum*, the species causing the most severe form of malaria in humans, PfHP1 is associated with the subtelomeric regions of all chromosomes and with some chromosome-internal islands (12, 30) where H3K9me3 is also enriched (35, 36). These heterochromatic regions contain over 400 protein-coding genes, most of which belong to gene families encoding exported virulence proteins and variant surface antigens including the *var* gene family (12, 30). The *var* gene family consists of 60 paralogs encoding antigenically and functionally distinct variants of *P. falciparum* erythrocyte membrane protein 1 (PfEMP1) that are displayed on the surface of infected red blood cells (iRBCs) (37–40). The interaction of PfEMP1 with receptors on endothelial cells or uninfected RBCs results in cellular adherence and sequestration of infected red blood cells (iRBCs) in the microvasculature, which is a major cause of severe malaria symptoms (41–43). In addition, antigenic variation and sequence diversity of PfEMP1 variants contribute significantly to immune evasion and hence to the establishment of chronic infection (41). Antigenic variation of PfEMP1 is based on switches in the mutually exclusive transcription of *var* genes (44). At any given time, only a single *var* gene is expressed while the remaining *var* gene family members are transcriptionally silenced (singular gene choice) (41, 44–46). *var* gene silencing is linked to the presence of H3K9me3/PfHP1 at the promoter and coding region (12, 13, 35, 47–50). The single active *var* gene, however, is instead enriched in H3K9ac and H3K4me3 in the upstream regulatory region (47). How the switch from the silenced to the active state is mediated is not known, but perinuclear locus reposition is linked to this process (51–56).

In addition to virulence gene families, HP1 also silences the gene encoding AP2-G, the master transcriptional regulator of gametocytogenesis in malaria parasites (29, 30, 49, 57–59). AP2-G is a member of the ApiAP2 family of putative transcription factors of apicomplexan parasites (60). The *ap2-g* gene is located in a chromosome-internal H3K9me3/HP1-demarcated heterochromatic island (12, 29, 30, 35, 36). Work in *P. falciparum* has shown that the PfHP1-dependent silencing of *pfap2-g* prevents sexual conversion and secures continuous parasite proliferation cycles. Removal of PfHP1 from the *pfap2-g* locus triggers expression of PfAP2-G and irreversible sexual conversion (49). Parasites that express AP2-G give rise to sexually committed progeny that exit the mitotic cell cycle and differentiate into female or male gametocytes (49, 57, 58, 61–65). Gametocytes are the only parasite stages capable of infecting the mosquito vector and as such are essential for malaria transmission. In *P. falciparum*, activation of *pfap2-g* transcription depends on the nuclear protein gametocyte development 1 (GDV1) that binds to and expels PfHP1 from the *pfap2-g* locus (61). Interestingly, even though *ap2-g* is associated with HP1 in all *Plasmodium* species (30), activation of this locus in *P. berghei* and other *Plasmodium* species infecting rodents is independent of GDV1 as they lack a GDV1 ortholog (61, 66, 67).

By analyzing a conditional PfHP1 knockdown mutant, we previously demonstrated that PfHP1 is essential for gene silencing and mitotic parasite proliferation (49). In that study, PfHP1-depleted parasites completed the current intra-erythrocytic multiplication cycle and gave rise to ring stage progeny, in which transcription of *var* genes and other heterochromatic gene families was highly augmented. Approximately 50% of this progeny underwent gametocytogenesis (25-fold increase compared to the isogenic control population) due to transcriptional derepression of the *pfap2-g* locus in the previous cell cycle. The other half of the PfHP1-depleted progeny represented asexual parasites that failed to enter schizogony due to defective genome replication (49). To interrogate the function of PfHP1 in more detail, we now conducted a functional analysis of PfHP1 domains by combining CRISPR/Cas9-mediated genome editing and the DiCre/LoxP system for conditional mutagenesis (68–70). We show that the C-terminal 76 residues encompassing the CSD (amino acids 191-266) are responsible for targeting PfHP1 to the nucleus. We further demonstrate that all three PfHP1 domains are required for heterochromatin formation, parasite proliferation and *pfap2-g* silencing and are therefore indispensable for proper PfHP1 function. Finally, we reveal that the HP1 hinge and CSD domains are functionally conserved between *P. falciparum* and the rodent malaria parasite *P. berghei*.

## Results

### Investigation of the roles of PfHP1 domains in PfHP1 localisation

To begin studying the functional contribution of individual PfHP1 domains, we engineered parasites that allow for the conditional expression of PfHP1 mutants based on the DiCre-loxP system (68, 69) using CRISPR-Cas9-based gene editing. In these parasites, the floxed endogenous *pfhp1* gene is excised upon rapamycin (RAP)-induced activation of the DiCre recombinase and replaced with a recodonised *pfhp1* gene encoding a mutated PfHP1 protein carrying a C-terminal GFP tag (Figures 1A and S1). We successfully used this approach recently to study the role of PfHP1 phosphorylation in regulating PfHP1 function (70). Here, we generated four such conditional PfHP1 mutant cell lines called 3D7/HP1-KO, 3D7/HP1-ΔCD, 3D7/HP1-ΔHinge, 3D7/HP1-ΔCSD, where amino acid residues 30-266 (full-length PfHP1), 30-58 (CD domain), 75-177 (hinge region) or 191-266 (CSD domain), respectively, are deleted upon RAP treatment (Figure 1B). In the HP1-ΔHinge mutant we substituted the hinge domain (102 amino acids) with a short linker peptide (22 amino acids) derived from the ApiAP2 transcription factor PfSIP2 where this peptide connects the two adjacent AP2 DNA-binding domains (71). The CRISPR-Cas9-based gene editing strategy used to generate these parasite lines is explained in detail in the Materials and Methods section and in Figure S1. PCR on parasite genomic DNA (gDNA) was performed to confirm the correct integration of the recodonised mutant *pfhp1-gfp* gene variants directly downstream of the endogenous *pfhp1* gene as well as the successful DiCre-mediated excision of the floxed endogenous *pfhp1* gene upon RAP treatment in all cell lines (Figure S1).

**Figure 1.**
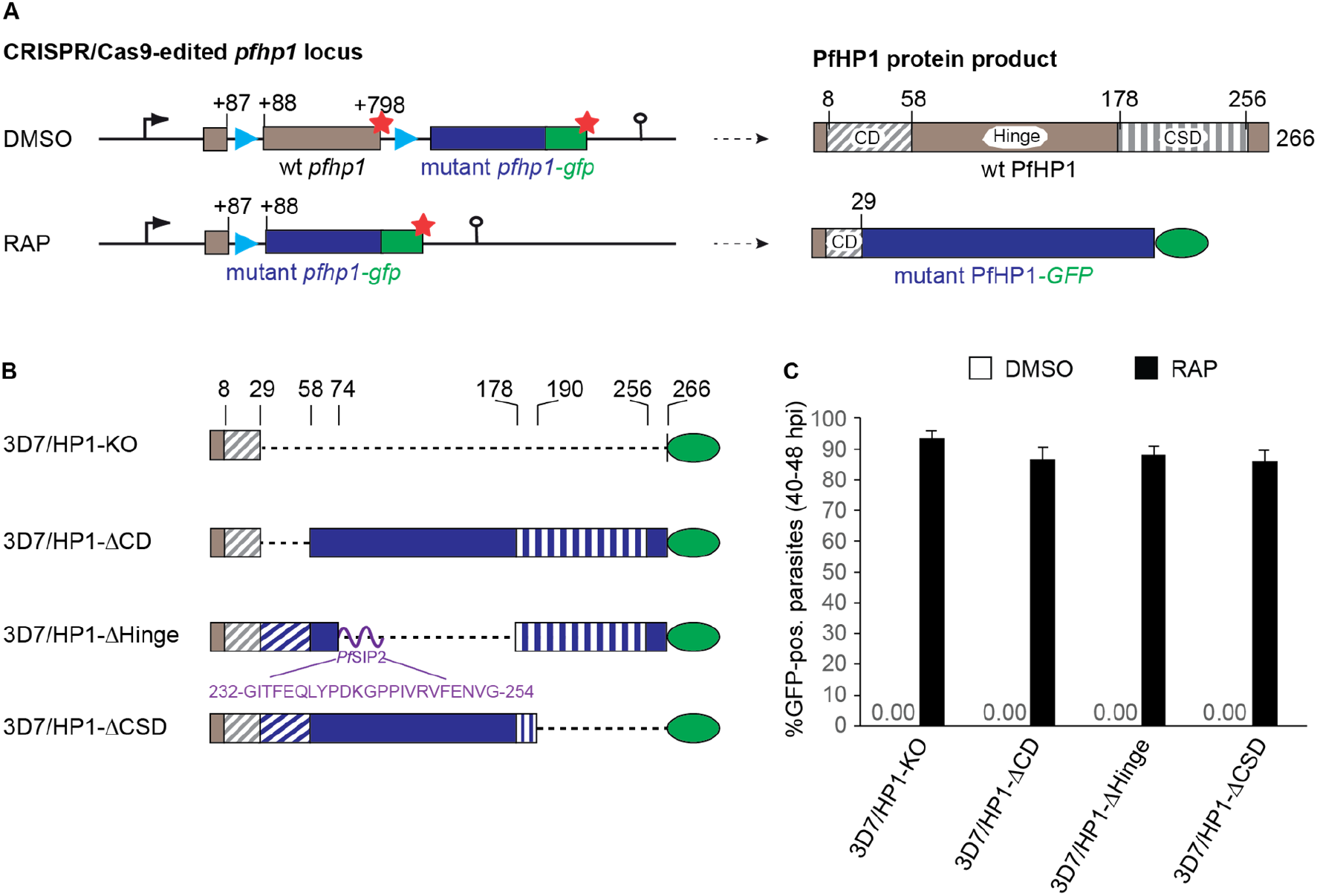
Generation of DiCre-inducible PfHP1 truncation mutants. **(A)** Schematics of the CRISPR/Cas9-edited *pfhp1* locus (left panel) and corresponding PfHP1 protein products (right panel) before (DMSO) and after (RAP) rapamycin-induced DiCre-dependent excision of the floxed wild type *pfhp1* locus. Blue arrowheads indicate the position of *sera2* intron:loxP elements. Red stars indicate STOP codons. Brown and blue boxes represent the wild type and the replacing mutant sequences *pfhp1/PfHP1*, respectively. Green boxes represent the *gfp/GFP* sequence. The CD, hinge and CSD domains within wild type PfHP1 are indicated. Numbers in the gene and protein schematics refer to nucleotide and amino acid positions, respectively. **(B)** Diagrams showing the organization of the PfHP1 truncation mutants expressed in the 3D7/HP1-KO, 3D7/HP1-ΔCD, 3D7/HP1-ΔHinge and 3D7/HP1-ΔCSD lines after RAP treatment. Dashed lines represent corresponding deletions in the mutant PfHP1 protein sequences. Brown and blue colours represent the remaining wild type PfHP1 N-terminus and the replacing mutant protein sequences, respectively. The CD and CSD are indicated by diagonal and vertical dashed stripes, respectively. The purple curved line represents the short PfSIP2-derived linker polypeptide between the CD and CSD domains in the PfHP1-ΔHinge mutant. The amino acid sequence of this linker and its position within the PfSIP2 protein are indicated. Numbers on top indicate amino acid positions within wild type PfHP1. **(C)** Proportion of GFP-positive parasites observed 40 hours after treatment with RAP or DMSO (control). Values represent the mean of three independent biological replicates (error bars indicate SD). For each sample >200 iRBCs were counted.

Scoring GFP-positive parasites by live cell fluorescence imaging at 40-48 hours post-invasion (hpi) (generation 1, 40 hrs post RAP treatment) confirmed the highly efficient excision of the endogenous *pfhp1* gene and expression of PfHP1-GFP mutant proteins upon RAP treatment in all four transgenic parasite lines (Figure 1C). In contrast, and as expected, parasites in the DMSO-treated control populations did not express the GFP-tagged PfHP1 mutant proteins (Figure 1C). With regard to subcellular localisation, we observed that the PfHP1 ΔCD-GFP and PfHP1 ΔHinge-GFP fusion proteins localized to the nucleus (Figure 2). The PfHP1ΔCSD-GFP protein exhibited reduced nuclear staining and a substantial fraction localised to the cytoplasm (Figure 2). Thus, the C-terminal polypeptide encompassing the CSD (amino acids 191 to 266) is required for efficient targeting of PfHP1 to the nucleus. Consistent with this observation the small fusion protein expressed in the 3D7/HP1-KO line, which only contains the first 29 amino acids of PfHP1 fused to GFP, localized to the cytoplasm (Figure 2B). The NucPred (72) and PSORTII (https://psort.hgc.jp/form2.html) algorithms identified putative canonical nuclear localisation signals (NLSs) in each of the three PfHP1 domains: a KKKK motif in the CD domain (amino acids 17 to 20), a PRRK motif in the hinge domain (amino acids 100 to 103) and a RRKK motif in the CSD domain (amino acids 201 to 204). Our fluorescence microscopy-based analysis suggest that the NLS consensus motif in the CSD domain may function as a major nuclear import element. The predicted NLS in the hinge domain may mediate some level of nuclear targeting whereas the KKKK motif in the CD seems to lack such activity. Importantly, none of the three PfHP1 domain deletion mutants assembled into heterochromatin domains but localised diffusely throughout the nucleoplasm. Typical perinuclear heterochromatic foci could only be observed in control parasites expressing full-length PfHP1-GFP after RAP treatment (3D7/HP1-Control) (Figure 2).

In summary, these data show that the C-terminal 76 amino acids comprising the CSD are responsible for the efficient targeting of PfHP1 to the nucleus. Inside the nucleus, the CD, hinge domain and CSD of PfHP1 are all required for the formation of perinuclear heterochromatin.

**Figure 2.**
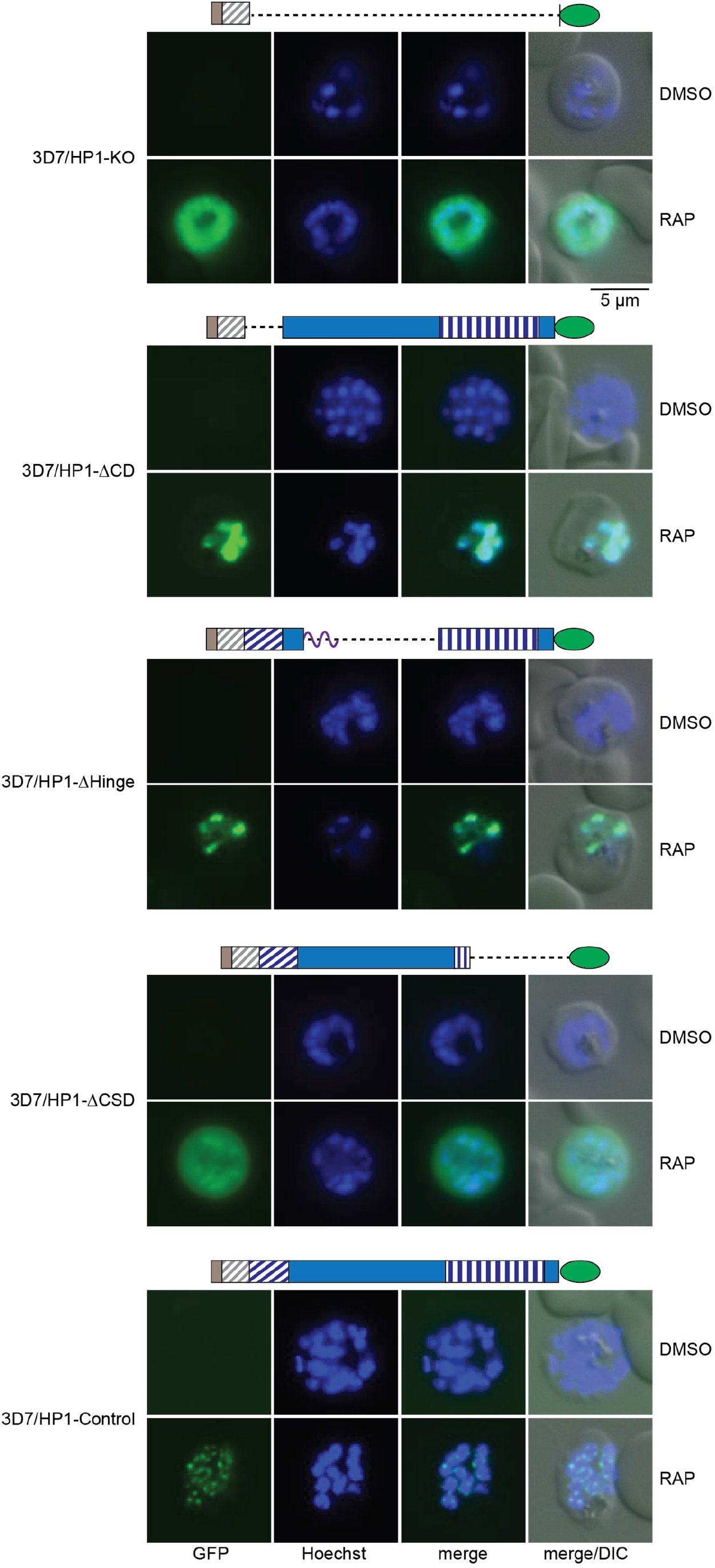
Subcellular localization of PfHP1 truncation mutants. Representative live cell fluorescence images showing the localization of the PfHP1-GFP-fusions in DMSO- and RAP-treated 3D7/HP1-KO, 3D7/HP1-ΔCD, 3D7/HP1-ΔHinge, 3D7/HP1-ΔCSD and 3D7/HP1-Control lines at late schizont stage (LS, 40-48 hpi, generation 1; 40 hrs after RAP treatment). Nuclei were stained with Hoechst. DIC, differential interference contrast. Scale bar, 5 μm.

### Each PfHP1 domains is essential for asexual proliferation of blood stage parasites

We next investigated the importance of each of the three PfHP1 domains for PfHP1 function. First, we compared parasite proliferation in the four PfHP1 truncation mutants grown under control conditions (DMSO) and after RAP treatment. To this end, parasite cultures (0.1% ring stage parasitaemia) were split (generation 1) and one half was treated with 100 nM RAP and the other half was treated with the DMSO solvent (control). For each cell line, the paired populations were followed over three generations by assessing the parasitaemia in the second and third generation by inspection of Giemsa-stained blood smears. As shown in Figure 3A, the parasitaemia of all DMSO-treated control populations increased 5-7-fold during each of the two multiplication cycles as expected. The RAP-treated 3D7/HP1-KO line showed a two-fold reduced parasite multiplication rate (PMR) after the first replication cycle and the progeny failed to replicate further (Figure 3A). This result is entirely consistent with our previous study using a conditional PfHP1-GFP-DD knockdown line, where PfHP1-depleted parasites multiplied at a two-fold reduced rate during the first cycle and were unable to proliferate further [Brancucci et al 2014]. Interestingly, we observed the same phenotype for all three PfHP1 domain deletion mutants (Figure 3A), which demonstrates that each individual PfHP1 domain (CD, hinge, CSD) is required for the proper function of PfHP1 in controlling mitotic proliferation of asexual blood stage parasites.

**Figure 3.**
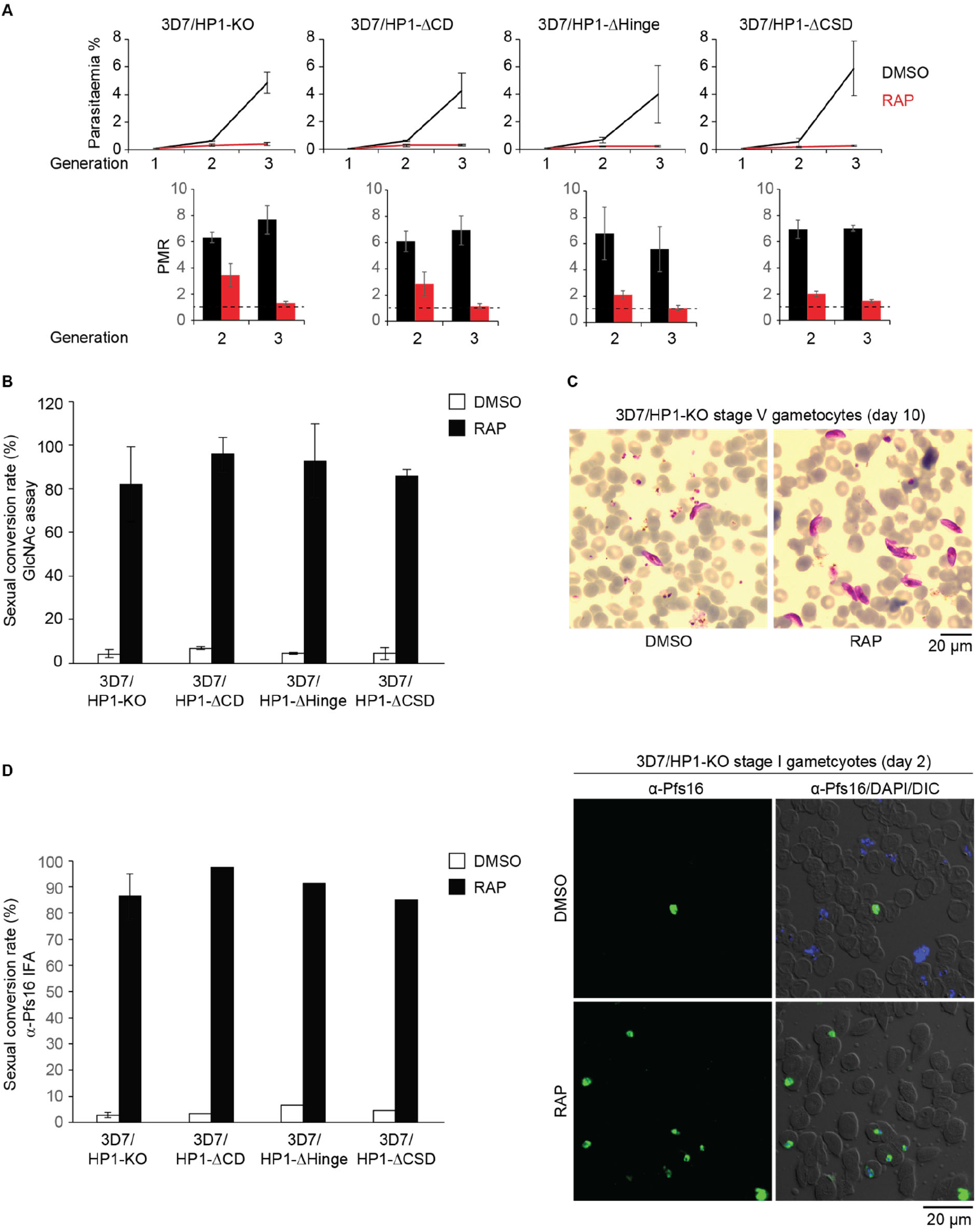
Phenotypes of PfHP1 truncation mutants. **(A)** Growth curves of the DMSO- and RAP-treated PfHP1 truncation mutants over three consecutive generations (top). Parasite multiplication rates (PMRs) reflect the fold increase in parasitaemia observed in generation 2 and 3 (bottom). Values are the mean of three biological replicates (error bars indicate SD). For each sample >3,000 RBCs were counted. **(B)** Sexual conversion rates of PfHP1 truncation mutants in DMSO- and RAP-treated parasites, assessed by inspection of Giemsa-stained blood smears of GlcNAc-treated cultures on day 6 of gametocytogenesis. Results are the mean of at least three replicates (error bars indicate SD). For each sample >3,000 RBCs were counted. **(C)** Representative overview images from Giemsa-stained blood smears showing stage V gametocytes (day 10 of gametocytogenesis) obtained from DMSO- and RAP-treated 3D7/HP1-KO parasites. Scale bar, 20 μm. **(D)** Sexual conversion rates of PfHP1 truncation mutants in DMSO- and RAP-treated parasites, assessed by α-Pfs16 IF As performed on stage I gametocytes on day 2 of gametocytogenesis. The result from the 3D7/HP1-KO line represents the mean of four replicates (error bar indicates SD). All other values derive from a single experiment. For each sample >200 iRBCs were counted. Representative overview images of α-Pfs16 IFAs used to quantify stage I gametocytes in DMSO- and RAP-treated populations of the 3D7/HP1-KO line are shown on the right. Scale bar, 20 μm.

We previously showed that next to the cell cycle-arrested trophozoites, the proliferation-defective progeny of the conditional PfHP1-GFP-DD knockdown mutant contained a large proportion of early stage gametocytes due to defective silencing of *pfap2-g* (49). Here, we examined a possible role of individual PfHP1 domains in controlling *pfap2-g* silencing by comparing sexual conversion rates (SCR) between DMSO- and RAP-treated parasites for all PfHP1 truncation mutants. To quantify SCRs, the ring stage progeny of DMSO- and RAP-treated parasites (generation 2, day 1 of gametocytogenesis) were cultured in medium containing N-acetyl-D-glucosamine (GlcNAc) to eliminate asexual parasites (73, 74) and the gametocytaemia determined after six days of GlcNAc treatment (stage III gametocytes). The DMSO-treated control populations of all conditional PfHP1 truncation mutants and the 3D7/HP1-Control line consistently displayed default SCRs of 4-7% (Figure 3B). As expected from the results previously obtained with the conditional PfHP1-GFP-DD knockdown line (49), the SCR of RAP-treated 3D7/HP1-KO parasites was massively increased reaching 82.1% (±17.3 SD), which is even higher compared to the average SCR of 52% reported upon knocking down PfHP1-GFP-DD expression (Figure 3B). Furthermore, similar to PfHP1-GFP-DD knockdown gametocytes, PfHP1 null gametocytes of the 3D7/HP1-KO line differentiated into stage V gametocytes in absence of PfHP1 expression (Figure 3C). Strikingly, we found that all three RAP-treated 3D7/HP1-ΔCD, 3D7/HP1-ΔHinge and 3D7/HP1-ΔCSD PfHP1 domain deletion mutants displayed similarly high SCRs of 85-95% (Figure 3B). These results were independently confirmed with immunofluorescence assays (IFAs) quantifying SCRs based on expression of the gametocyte-specific marker Pfs16 processes (75) in the progeny of DMSO- and RAP-treated parasites (40-48hpi, day 2 of gametoyctogenesis) (Figure 3D). Together, these findings demonstrate that all three PfHP1 domains are essential for the function of PfHP1 in mediating gene silencing and reinforce the central role for PfHP1 in suppressing sexual conversion.

### The function of the HP1 hinge and chromo shadow domains are conserved between *P. falciparum* and *P. berghei*

It has been shown that replacement of the CD domain of *S. pombe* Swi6 with the CD domain of the mouse HP1 protein M31 retained Swi6 function in sporulation, normal zygote asci formation and mitotic stability. CSD substitution, however, did not restore Swi6 function in these processes (26). Here, we tested the functional conservation between the HP1 orthologs of *P. falciparum* and *P. berghei*, the most widely used mouse malaria model parasite. We were primarily interested in this question because *P. berghei* parasites lack the GDV1 protein that is required for PfHP1 eviction from the *pfap2-g* locus and subsequent sexual conversion in *P. falciparum* (61, 66).

PfHP1 (266 amino acids) and PbHP1 (281 amino acids) display an overall sequence identity of 68%. The CDs and CSDs are highly conserved with 88% and 90% identical amino acids, respectively, whereas the intervening sequence encompassing the hinge domain shows poor conservation with only 45% sequence identity (Figure S2). We first attempted to obtain a transgenic line where RAP treatment would replace *pfhp1* with the *pbhp1* gene. For unknown reasons, however, we repeatedly failed to integrate the corresponding conditional expression cassette into the endogenous *pfhp1* locus. Hence, we decided to perform domain swap experiments and generated two DiCre-inducible PfHP1-hybrid cell lines, namely 3D7/HP1-hyb-PbHinge and 3D7/HP1-hyb-PbCSD. In these cell lines, RAP treatment leads to the replacement of the PfHP1 hinge or CSD domain with the PbHP1 hinge or CSD domain, respectively, and both hybrid HP1 proteins are expressed as C-terminal GFP fusions (Figure 4A and Figure S1). Note that a CD domain swap cell line could not be generated for technical reasons as the 3D7/N31DC mother line used to generate the conditional mutants does not allow to swap the entire CD domain (Figure S1). PCR on gDNA confirmed the correct insertion of the two hybrid *hp1* sequences downstream of the endogenous *pfhp1* gene and the successful replacement of endogenous *pfhp1* with the hybrid genes after RAP-induced DiCre-mediated recombination in both lines (Figure S1). To assess the ability of the HP1 hybrid proteins in forming heterochromatic domains, we performed live cell fluorescence imaging at 40-48 hpi (generation 1; 40 hrs post RAP treatment) and at 16-24 hpi in generation 2 ring stages to observe the localization of the GFP fusion proteins. As expected, no GFP signal was detectable in both DMSO-treated control parasites (Figures 4B and 4C). After RAP treatment, over 95% of parasites expressed the GFP-tagged hybrid HP1 proteins (Figure 4B) and both PfHP1-hyb-PbHinge-GFP and PfHP1-hyb-PbCSD-GFP localized to perinuclear heterochromatin foci in schizonts and the ring stage progeny in a pattern indistinguishable from that observed for wild type PfHP1-GFP (49, 61, 70) (Figures 2 and 4C). Thus, replacement of the PfHP1 hinge or CSD domain with those from PbHP1 retains proper heterochromatin localization of HP1 in *P. falciparum*.

**Figure 4.**
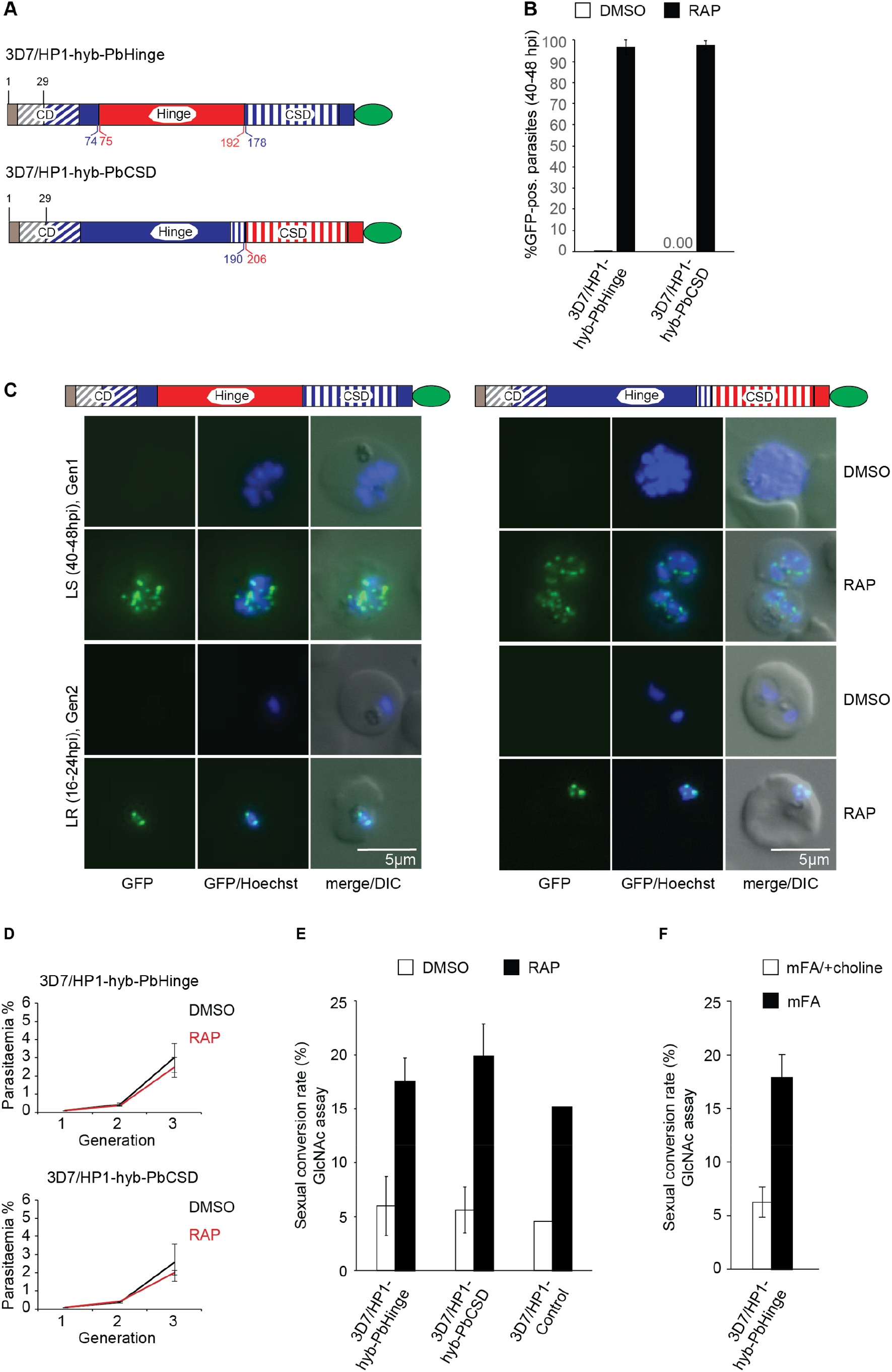
Generation of DiCre-inducible PfHP1-PbHP1 hybrid mutants. **(A)** Diagrams showing the GFP-tagged PfHP1-PbHP1 hybrid proteins expressed in the 3D7/HP1-hyb-PbHinge and 3D7/HP1-hyb-PbCSD cell lines after RAP treatment. Brown and blue colours represent the remaining wild type PfHP1 N-terminus and the replacing PfHP1 protein sequences, respectively. Red colours identify the hinge domain and CSD derived from PbHP1. The CD and CSD are indicated by diagonal and vertical dashed stripes, respectively. Numbers in blue and red refer to amino acid positions within the PfHP1 and PbHP1 sequence, respectively. **(B)** Proportion of GFP-positive parasites observed 40 hours after treatment with RAP or DMSO (control). Values represent the mean of three independent biological replicates (error bars indicate SD). For each sample >140 iRBCs were counted. **(C)** Representative live cell fluorescence images showing the localization of GFP-tagged PfHP1-PbHP1 hybrid proteins in 3D7/HP1-hyb-PbHinge and 3D7/HP1-hyb-PbCSD parasites in late schizonts (LS, 40-48 hpi, generation 1, 40 hrs after RAP treatment) and in the late ring stage progeny (LR, 16-24 hpi, generation 2). Nuclei were stained with Hoechst. DIC, differential interference contrast. Scale bar, 5 μm. **(D)** Growth curves of the DMSO- and RAP-treated 3D7/HP1-hyb-PbHinge and 3D7/HP1-hyb-PbCSD parasites over three consecutive generations. Values are the mean of at least three independent replicate experiments (error bars represent SD). For each sample >3,000 RBCs were counted. **(E)** Sexual conversion rates of the DMSO- and RAP-treated 3D7/HP1-hyb-PbHinge and 3D7/HP1-hyb-PbCSD mutants and the 3D7/HP1-Control line, assessed by inspection of Giemsa-stained blood smears of GlcNAc-treated cultures on day 6 of gametocytogenesis. Values represent the mean of at least three independent replicate experiments (error bars represent SD). The values for the 3D7/HP1-Control line derive from a single experiment and are consistent with previously published data (70). For each sample >3,000 RBCs were counted. **(F)** Sexual conversion rates of 3D7/HP1-hyb-PbHinge-expressing parasites cultured in minimal fatty acid medium (mFA) or mFA supplemented with 2 mM choline (mFA/+choline), assessed by inspection of Giemsa-stained blood smears of GlcNAc-treated cultures on day 6 of gametocytogenesis. Values represent the mean of two independent replicate experiments (error bars represent SD). For each sample >1,800 RBCs were counted.

We next evaluated parasite proliferation in the 3D7/HP1-hyb-PbHinge and 3D7/HP1-hyb-PbCSD lines over three generations and observed no major differences in parasite multiplication between DMSO- and RAP-treated parasites, revealing proper function of the HP1 hybrid proteins in controlling mitotic cell cycle progression and parasite proliferation (Figure 4D). This observation also indicated that the HP1 hybrid proteins mediate efficient silencing of the *pfap2-g* locus. To confirm this property, we compared the SCRs of DMSO- and RAP-treated parasites for both HP1 hybrid lines and the 3D7/HP1-Control line using the GlcNAc assays as described above. As shown in Figure 4E, the RAP-treated 3D7/HP1-hyb-PbHinge and 3D7/HP1-hyb-PbCSD populations showed a 3-4-fold increased SCR compared to the DMSO-treated controls. However, RAP-treated 3D7/HP1-Control parasites also showed a 3-4-fold increased SCR in the RAP treatment cycle, which has been observed previously and is caused by unknown mechanisms linked to the DiCre-mediated exchange of endogenous *pfhp1* with the recodonised *pfhp1-gfp* gene (70). Hence, the PfHP1-hyb-PbHinge and PfHP1-hyb-PbCSD proteins appear to silence the *pfap2-g* locus as efficiently as wild type PfHP1. Furthermore, since 3D7/HP1-hyb-PbHinge and 3D7/HP1-hyb-PbCSD parasites were still able to produce gametocytes at a rate similar to the 3D7/HP1-Control line, it seems that GDV1 can interact with and evict the hybrid HP1 proteins from the *pfap2-g* locus. To confirm this hypothesis, we induced sexual commitment in 3D7/HP1-hyb-PbHinge parasites using minimal fatty acid (mFA) medium lacking lysophosphatidylcholine (lysoPC) and choline (67). While the exact mechanisms involved in this sensing pathway remain elusive, the increased sexual commitment rates observed under lysoPC/choline-depleted conditions correlates with an increased proportion of parasites expressing GDV1 (61). To this end, 3D7/HP1-hyb-PbHinge parasites were treated with RAP to swap endogenous *pfhp1* with the *pfhp1-hyb-pbhinge* hybrid gene. After one additional multiplication cycle, PfHP1-hyb-PbHinge-expressing parasites were split, cultured separately in mFA (induces sexual commitment) or mFA supplemented with 2 mM choline (mFA/+choline) (suppresses sexual commitment) (67) and SCRs were determined in the progeny using the GlcNAc assay. As shown in Figure 4F, 3D7/HP1-hyb-PbHinge parasites were indeed responsive to choline depletion, showing 3-fold increased SCRs compared to the same population grown in presence of choline.

In summary, even though we failed to generate a *P. falciparum* line expressing full-length PbHP1, our findings obtained with the hinge and CSD domain swap mutants demonstrate that PbHP1 can execute the core functions of PfHP1 in proliferation and gene silencing in *P. falciparum* parasites. They furthermore suggest that PbHP1 can still functionally interact with GDV1 even though *P. berghei* parasites do not possess a GDV1 ortholog.

## Discussion

Here, we used a DiCre-based conditional protein domain deletion and swapping approach to study the function of PfHP1 in *P. falciparum* blood stage parasites. By combining the data gained from subcellular protein localisation analysis and parasite multiplication and sexual differentiation assays, we are able to draw important conclusions as to the specific roles each domain plays in mediating proper PfHP1 function.

Previous studies indicated that in different eukaryotes different domains of HP1 carry the sequence information required for targeting HP1 to the nucleus and that these regions do not always possess canonical NLS sequences (19, 26–28, 76, 77). Therefore, we could not rely on sequence homology to predict the nuclear targeting sequences within PfHP1 but had to determine them experimentally instead. We found that the C-terminal 76 residues encoding the CSD (amino acids 191-266) are essential for effective nuclear targeting as only the PfHP1-ΔCD and PfHP1-ΔHinge fusions, but not the PfHP1-ΔCSD fusion, were efficiently imported into the nucleus. It is possible that the predicted NLS motif in the CSD (RRKK; amino acids 201 to 204) is indeed directly responsible for directing PfHP1 into the nucleus. The predicted NLS element in the hinge domain (PRRK; amino acids 100 to 103) may also allow some level of nuclear import as a fraction of PfHP1-ΔCSD proteins still localized to the nucleus. However, since the bioinformatic identification of NLSs in *P. falciparum* proteins has only weak predictive power (78), mutational analyses of these putative NLS motifs will be required to test whether and to what extent they are implicated in mediating nuclear import of PfHP1.

Interestingly, none of the three PfHP1 domain deletion mutants was capable of forming and inheriting heterochromatin to daughter nuclei during schizogony. Heterochromatin binding of HP1 has been well investigated in many model organisms (19, 26–28, 77). Studies employing different experimental systems highlighted the contribution of more than one structural HP1 domain for proper heterochromatin targeting. In mice, for instance, the heterochromatin-targeting ability of HP1α involves RNA binding via a region in the hinge domain together with the binding to methylated histone via the CD (19). In *D. melanogaster*, the domains targeting HP1 to the nucleus and to heterochromatin were identified by analysing a panel of HP1 truncation mutants tagged with β-galactosidase at the N-terminus (27). Similar to our study, protein fusions containing the CD and/or hinge domain but lacking the CSD failed to localise to the nucleus, while protein fusions containing the majority of the CSD region (amino acids 152-206; CSD encompasses amino acids 142-206) showed nuclear localization. However, proper assembly into heterochromatin was only found for protein fusions containing a substantial part of the hinge domain and the CSD region (amino acids 95-206) (27). Given that in these experiments the β-galactosidase-HP1 fusions were ectopically expressed in a wild-type HP1 background, successful heterochromatin targeting may have resulted from hetero-dimerization between CSD-containing fusion proteins and endogenous HP1. In our study, however, the PfHP1-GFP truncation mutants were expressed in a PfHP1 null background and only full length PfHP1 was able to assemble into perinuclear heterochromatic foci during schizogony. To interpret these findings it is useful to bring to mind the process of parasite multiplication by schizogony, where four to five subsequent rounds of genome duplication and nuclear division precede daughter cell formation. The re-establishment of heterochromatin on newly replicated chromosomes is likely initiated by the binding of PfHP1 to existing H3K9me3-containing nucleosomes that are distributed between the replicated chromosomes in a semi-conservative manner. Heterochromatin spreading, however, will require *de novo* methylation of H3K9 on newly incorporated nucleosomes by a H3K9-specific SUVAR3-9-like HKMT [presumably PfSET3 (36, 79, 80)], the recruitment of which depends on its interaction with the CSD of HP1 (5, 17). Hence, mutations that prevent PfHP1 either from binding to chromatin or from recruiting the H3K9-specific HKMT will equally result in defective heterochromatin formation and this defect will become more pronounced with each additional round of genome duplication during schizogony. The fact that the PfHP1 ΔCD-GFP, PfHP1 ΔHinge-GFP and PfHP1 ΔCSD-GFP deletion mutants all failed to assemble into perinuclear heterochromatic foci in schizonts and the subsequent ring stage progeny shows that each of the three domains is essential for the *de novo* heterochromatin assembly on replicated chromosomes. While this observation is not surprising with regard to the CD (required for binding to H3K9me3) and CSD (required for PfHP1 dimerisation and HKMT recruitment), our results demonstrate that the PfHP1 hinge domain is also indispensable for proper heterochromatin assembly. Studies on mouse and *Xenopus laevis* HP1 proteins have shown that the RNA- and DNA-binding capacity of the hinge domain is important for high affinity binding of HP1 to H3K9me3 and stable heterochromatin formation (18–20). We therefore anticipate the crucial role of the PfHP1 hinge domain in heterochromatin assembly may be based on similar properties.

Conditional expression of the three PfHP1 domain deletion mutants phenocopied the conditional knockout of PfHP1 in the 3D7/HP1-KO line. All four mutants produced heterochromatin-depleted ring stage progeny that consisted of up to 95% sexually committed parasites and a small proportion of asexual parasites that failed to proliferate further. This phenotype is identical but more pronounced compared to that obtained upon knocking down PfHP1-GFP-DD expression (49). In their previous study, Brancucci and colleagues demonstrated that conditional depletion of PfHP1 expression released the *pfap2-g* locus from PfHP1-dependent silencing, which in turn triggered PfAP2-G expression and sexual conversion in 52% of the ring stage progeny. The other half of the progeny represented asexual parasites that arrested at the trophozoite stage due to failed entry into S-phase (49). The substantially higher proportion of sexually committed ring stage progeny observed in the inducible PfHP1 KO or domain deletion mutants generated here shows that the DiCre-mediated excision or mutagenesis of the *pfhp1* gene leads to an almost complete abolishment of PfHP1-dependent gene silencing. Furthermore, these results also demonstrate that the lack of perinculear heterochromatic foci in the PfHP1 domain deletion mutants is also reflected at the functional level in defective gene silencing.

Previous chromatin immunoprecipitation followed by next generation sequencing (ChIP-seq) experiments performed on six different *Plasmodium* species (*P. falciparum*, *P. vivax*, *P. knowlesi*, *P. berghei, P. yoelii, P. chabaudi)* suggest a conserved role for heterochromatin in facilitating the clonally variant expression of species-, clade- and genus-specific genes that are primarily involved in hostparasite interactions during blood stage infection (29, 30). In addition, HP1 also controls the heritable silencing of *ap2-g* in all malaria parasite species examined to date (29, 30). In *P. falciparum*, activation of *pfap2-g* expression and subsequent sexual conversion is triggered by the GDV1-dependent displacement of PfHP1 from the *pfap2-g* locus (61). Notably, GDV1 is essential for this process because GDV1 loss-of-function mutants are unable to commit to gametocytogenesis (63, 66, 81). Expression of GDV1 itself appears to be negatively regulated by a long non-coding antisense RNA (61, 82) and low concentrations of the serum lipid lysoPC induce GDV1 expression through an unknown sensing pathway (61). In addition to *P. falciparum*, all other *Plasmodium* spp. infecting primates as well as avian malaria parasites (*P. gallinaceum, P. relictum)* possess a GDV1 ortholog (www.plasmodb.org), suggesting these species may employ a conserved strategy to activate *ap2-g* expression in response to environmental stimuli. Intriguingly, however, *Plasmodium* spp. infecting rodents lost the *gdv1* locus (61, 66, 67) and sexual commitment in *P. berghei* is insensitive to lysoPC depletion (67). Whether *ap2-g* activation in these parasites requires an unknown factor functionally equivalent to GDV1 and responsive to an alternative sensing pathway, or whether PbHP1 is less stably associated with the *pbap2-g* locus allowing for a higher level of stochastic activation is unknown at this stage. Our results show that the hinge domain and the CSD of PbHP1 can fully complement PfHP1’s function in mitotic proliferation, heterochromatin formation and *pfap2-g* silencing. Because 3D7/HP1-hyb-PbHinge and 3D7/HP1-hyb-PbCSD parasites did not display higher default SCRs than the 3D7/HP1-Control line, both hybrid HP1 proteins seem to form similarly stable heterochromatin domains as compared to wild type PfHP1, at least in *P. falciparum*. Importantly, our findings also demonstrate that GDV1 can still recognise and displace both HP1 hybrid proteins from the *pfap2-g* locus. Although the physical interaction between GDV1 and PfHP1 has not yet been mapped to a particular PfHP1 domain (REF), we speculate that GDV1 may interact with the CSD for two reasons. First, the CSD dimer interface is responsible for most interactions between HP1 and other regulatory factors in model eukaryotes (7–9, 11). Second, in contrast to their hinge domains the CSDs of PfHP1 and PbHP1 share high sequence identity and perfect conservation of residues predicted to be involved in CSD homo-dimerization and protein-protein interaction (Figure S2). However, we cannot rule out the possibility that the N-terminus and/or the CD domain are important for this interaction. Irrespective of this uncertainty, our findings strongly suggest that PbHP1 retained the capacity to interact with GDV1 and that this interaction evolved based on evolutionary conserved features of HP1 orthologs in malaria parasites. In light of this conclusion, it may be interesting to explore if ectopic expression of GDV1 could be used as an experimental tool for the induction of high sexual conversion rates in *P. berghei*, as recently reported for *P. falciparum* (59, 61).

In summary, we demonstrate that each of the three PfHP1 domains is essential for heterochromatin formation, gene silencing and mitotic proliferation in *P. falciparum* blood stage parasites. We also discovered that the hinge domain and CSD of *P. berghei* HP1 fully complement the function of the corresponding domains of PfHP1 in these processes and sustain the capacity for GDV1-dependent eviction of HP1 from the *pfap2-g* locus. Together, these findings provide major new insight into HP1 function in malaria parasites and offer new possibilities for a further functional dissection of this essential silencing factor in parasite biology.

## Methods

### Parasite culture and transfection

The transgenic cell lines generated in this study were cultured at 5% hematocrit in RPMI-1640 medium supplemented with 25 mM HEPES, 100 mM hypoxanthine, 24 mM sodium bicarbonate, 0.5% Albumax II supplemented with 2mM choline to reduce default sexual conversion rates (SCR) as demonstrated recently (67). Parasite cultures were synchronized using 5% sorbitol as described previously (83). Cotransfection of CRISPR/Cas9 and donor plasmids into the DiCre-expressing line 3D7/1G5DiCre (69) and selection of transfected populations was performed as described recently (61, 70).

### Transfection constructs

We applied CRISPR/Cas9-mediated genome editing and the DiCre/LoxP system (68, 69) to generate parasite lines conditionally expressing truncated PfHP1 and hybrid HP1 variants. We engineered (1) 3D7/HP1-KO for full-length PfHP1 (PF3D7_1220900) deletion (deletion of amino acids 30-266); (2) 3D7/HP1-ΔCD for expression of a PfHP1 CD deletion mutant (deletion of amino acids 30-58); (3) 3D7/HP1-ΔHinge for expression of a PfHP1 hinge domain deletion mutant (amino acids 75-177 of PfHP1 were replaced with a linker peptide representing amino acid 232-254 of PfSIP2 (PF3D7_0604100)); (4) 3D7/HP1-ΔCSD for expression of a PfHP1 CSD deletion mutant (deletion of amino acids 191-266); (5) 3D7/HP1-hyb-PbHinge for expression of a chimeric PfHP1, in which the PfHP1 hinge domain (amino acids 75-177) was replaced by the PbHP1 (PBANKA_1436100) hinge domain (amino acids 75-192); and (6) 3D7/HP1-hyb-PbCSD for expression of a chimeric PfHP1, in which the PfHP1 CSD domain (amino acids 191-266) was replaced by the PbHP1 CSD domain (amino acid 206-281). To obtain these cell lines, we transfected the mother cell line 3D7-1G5DC/5’-loxPint-g31 (or 3D7/N31DC), which carries a *sera2* intron:loxP element (68) within the 5’ end of the *pfhp1* coding sequence (70), with donor plasmids to insert a second *sera2* intron:loxP sequence directly downstream of the endogenous *pfhp1* stop codon, followed by either the *gfp* coding sequence (HP1-KO), or a recodonised mutated *pfhp1* sequence fused to *gfp* (PfHP1-ΔCD, PfHP1-ΔHinge, PfHP1-ΔCSD, PfHP1-hyb-PbHinge, PfHP1-hyb-PbCSD). The donor plasmids are derivatives of the pD-PfHP1-Control donor plasmid described recently (70). Each donor plasmid was co-transfected with the CRISPR/Cas9 plasmid pBF-gC-guide250 that expresses SpCas9, a guide RNA targeting the 3’ end of the *pfhp1* coding sequence and the positive-negative drug selection marker blasticidin deaminase (BSD) fused to yeast cytosine deaminase/uridyl phosphoribosyl transferase (yFCU) (70).

Cloning of the pD-PfHP1-KO plasmid has recently been described (Bui et al.,) and consists of the pUC19 backbone carrying a 5’ homology region (HR) spanning bps +88 to +798 of the *pfhp1* gene terminating with a stop codon, followed by the 103 bp sera2 intron:loxP element (68), the *gfp* coding sequence ending with a stop codon, and a 3’ HR encompassing the 824 bps directly downstream of the *pfhp1* coding sequence. All additional donor plasmids described below carry the same 5’ and 3’ HR for homology-directed repair of the Cas9-induced DNA lesion.

The pD-PfHP1-ΔCD donor plasmid was constructed by Gibson assembly joining three PCR fragments encoding (1) the pD plasmid backbone amplified from pUC19 using primers PCRA_F and PCRA_R; (2) the *pfhp1* 5’ HR followed by the 103 bp *sera2* intron:loxP sequence amplified from pD-PfHP1-Control (70) using primers F158 and R143; and (3) a fragment amplified from pD-PfHP1-Control using primers F177 and R163 spanning, in the following order, bps +175 to +798 of a synthetic recodonized *pfhp1* coding sequence (70) omitting the stop codon, the *gfp* coding sequence ending with a stop codon and the *pfhp1* 3’ HR.

The pD-PfHP1-ΔHinge donor plasmid was constructed by Gibson assembly joining four PCR fragments encoding (1) the pD plasmid backbone amplified from pUC19 using primers PCRA_F and PCRA_R; (2) the *pfhp1* 5’ HR followed by the *sera2* intron: loxP sequence amplified from pD-PfHP1-Control using primers F158 and R143; (3) a fragment amplified from the pBcam-ΔHinge-3HA-Cherry plasmid (Supplementary Methods) using primers F164 and R165 spanning, in the following order, bps +88 to +222 of the recodonised *pfhp1* sequence, bps +694 to +762 of the *pfsip2* coding sequence encoding the linker region separating the tandem AP2 domains (71) and bps +532 to +798 of the recodonised *pfhp1* coding sequence omitting the stop codon; and (4) a fragment amplified from pFdon-C-loxP-g250 (70) using primers F162 and R163 spanning the *gfp* coding sequence and *pfhp1* 3’HR.

The pD-PfHP1-ΔCSD donor plasmid was constructed by Gibson assembly joining three PCR fragments encoding (1) the pD plasmid backbone amplified from pUC19 using primers PCRA_F and PCRA_R; (2) a fragment amplified from pD-PfHP1-Control using primers F158 and R178 spanning, in the following order, the *pfhp1* 5’HR followed by the *sera2* intron:loxP element and bps +88 to +570 of the recodonised *pfhp1* coding sequence; and (3) a fragment amplified from pFdon-C-loxP-g250 (70) using primers F162 and R163 spanning the *gfp* coding sequence and *pfhp1* 3’HR.

The pD-hyb-PbHinge donor plasmid was constructed by Gibson assembly joining two PCR fragments encoding (1) part of the pD-HP1-KO plasmid amplified from its own template using primers F162 and R143 and representing, in the following order, the *gfp* coding sequence, the *pfhp1* 3’HR, the pUC19 plasmid backbone and the *pfhp1* 5’HR followed by the 103 bp *sera2* intron:loxP sequence; and (2) a fragment amplified from the pBcam-hyb-PbHinge-3HA-Cherry plasmid (Supplementary Methods) using primers F164 and R165 spanning, in the following order, bps +88 to +222 of the recodonised *pfhp1* coding sequence, bps +223 to +576 of the *pbhp1* coding sequence and bps +532 to +798 of the recodonised *pfhp1* coding sequence omitting the stop codon.

The pD-hyb-PbCSD donor plasmid was constructed by Gibson assembly as explained above for pD_hyb_Pbhinge joining two PCR fragments encoding (1) the corresponding part of the pD-HP1-KO plasmid backbone amplified from its own template using primers F162 and R143; and (2) a fragment amplified from the pBcam-hyb-PbCSD-3HA-Cherry (Supplementary Methods) using primers F164 and R161 spanning bps +88 to +570 of the recodonised *pfhp1* coding sequence followed by bps +616 to +843 of the *pbhp1* sequence omitting the stop codon.

For each transfection, 50 μg of the pBF-gC-guide250 plasmid was mixed with 50 μg of donor plasmid and electroporated into the 3D7/N31DC mother parasite line. Transfected parasites were selected on 5 μg/ml BSD-S-HCl and established as described previously (61). All oligonucleotide sequences used for the cloning of donor plasmids are provided in Supplementary Table S1.

### Induction of DiCre recombinase-mediated DNA excision by rapamycin treatment

Parasites were synchronized twice 16 hours apart to obtain an eight-hour growth window (16-24 hpi). After invasion into new RBCs, parasites were synchronized again at 0-8 hpi (generation 1) and split into two equal populations, one of which was treated with 0.02% DMSO (negative control) and the other half was treated with 100 nM RAP for 60-90 minutes as described previously (84). The cultures were then spun down, washed once and resuspended in culture medium lacking RAP for onward *in vitro* culture.

### Live cell fluorescence imaging and indirect immunofluorescence assays

To quantify the efficiency of *pfhp1* excision after RAP treatment, live cell fluorescence microscopy was performed as described before (85) with minor modification using 5 μg/ml Hoechst (Merck) to stain the nuclei. Excision efficiency was determined as the percentage of GFP-positive schizonts at 40-48 hpi in generation 1 (40 hours post RAP treatment) (>200 schizonts counted per experiment). IFAs were performed on methanol-fixed cells using mouse mAb α-Pfs16 (kind gift from Robert W. Sauerwein), 1:250; and Alexa Fluor 488-conjugated α-mouse IgG (Molecular Probes), 1:250. Nuclei were stained with 5 μg/ml Hoechst (Merck). Images were taken at 630-fold magnification on a Leica DM 5000B microscope with a Leica DFC 300 FX camera, acquired via the Leica IM1000 software and processed using ImageJ [https://imagej.nih.gov/ij/]. For each experiment, images were acquired and processed with identical settings.

### Parasite multiplication assay

Parasites were tightly synchronized twice 16 hours apart within the same intra-erythrocytic cycle. After invasion into new RBCs, parasites were split at 0-8 hpi (generation 1) into two equal populations, of which one half was treated with 0.02% DMSO (negative control) and the other half was induced for DiCre recombinase-mediated DNA excision by RAP treatment as described above. Giemsa smears were prepared to determine the parasitaemia at 16-24 hpi (generation 1). Giemsa-stained smears were also prepared two days later (generation 2) and four days later (generation 3). Parasitaemia was counted by visual inspection of Giemsa-stained blood smears (≥3’000 RBCs counted per experiment). Parasite multiplication rates (PMRs) were determined as the parasitaemia observed in the following generation divided by the parasitaemia observed in the previous generation.

### Gametocyte conversion assay

After DMSO or RAP treatment in generation 1, parasites were allowed to complete schizogony and re-invade RBCs. At 16-24 hpi in generation 2 (day 1 of gametocytogenesis), each pair of parasites was treated with 50 mM N-acetyl-D-glucosamine (GlcNAc) for six days to eliminate asexual parasites (73, 74) and then cultured with normal culture medium for another six days to observe the maturation of gametocytes. Gametocytaemia was determined on day 6 of GlcNAc treatment by visual inspection of Giemsa-stained blood smears. SCRs were determined as the gametocytaemia observed on day 6 of gametocyte development as a proportion of the total parasitaemia observed on day 1 of gametocytogenesis. For the experiment presented in Figure 4F, 3D7/HP1-hyb-PbHinge ring stage parasites (generation 1) were treated with RAP to swap endogenous *pfhp1* with the *pfhp1-hyb-pbhinge* hybrid gene and allowed to replicate and invade new RBCs. At 24-30 hpi (generation 2) the population was split and cultured separately in minimal fatty acid medium (mFA) or mFA supplemented with 2 mM choline (mFA/+choline) to induce or suppress sexual commitment, respectively, as previously described (61, 67). At 16-24 hpi in generation 3 (day 1 of gametocytogenesis) the paired populations were cultured in presence of 50 mM N-acetyl-D-glucosamine (GlcNAc) until day 6 of gametocyte development and SCRs were determined as described above.

### Genomic DNA isolation, polymerase chain reaction and Sanger sequencing

To evaluate correct editing of the *pfhp1* locus, PCRs on gDNA isolated from all transgenic parasite lines were performed. gDNAs were sampled and isolated as described previously (85). To evaluate the DNA excision efficiency after RAP treatment, PCRs were performed on gDNA isolated 24-36 hours post treatment. Primers were designed to allow PCR amplification across the 5’ to 3’ homology regions. All transfection plasmids generated in this study have been validated by Sanger sequencing. All transfection plasmids have been designed and Sanger sequencing results analysed using the SnapGene software (GSL Biotech; www.snapgene.com). All PCR primer sequences are listed in Supplementary Table S1.

### Data availability

All data generated or analysed during this study are included in this published article and its Supplementary Information files.

## Acknowledgements

This work has been supported by the Swiss National Science Foundation (grant numbers 31003A_143916 and 31003A_163258).

## Author contributions

H.T.N.B. designed and performed experiments, analysed data, prepared illustrations and wrote the manuscript. A.P. helped performing sexual conversion assays and N.M.B.B. supervised these experiments and provided resources and conceptual advice. T.S.V. conceived of the study, designed, supervised, and analysed experiments, prepared illustrations, provided resources and wrote the manuscript. All authors contributed to editing of the manuscript.

## References

1. James TC, Elgin SC. 1986. Identification of a nonhistone chromosomal protein associated with heterochromatin in Drosophila melanogaster and its gene. Mol Cell Biol 6:3862–3872.

2. Eissenberg JC, James TC, Foster-Hartnett DM, Hartnett T, Ngan V, Elgin SC. 1990. Mutation in a heterochromatin-specific chromosomal protein is associated with suppression of position-effect variegation in Drosophila melanogaster. Proc Natl Acad Sci U S A 87:9923–9927.

3. Lachner M, O’Carroll D, Rea S, Mechtler K, Jenuwein T. 2001. Methylation of histone H3 lysine 9 creates a binding site for HP1 proteins. Nature 410:116–120.

4. Bannister AJ, Zegerman P, Partridge JF, Miska EA, Thomas JO, Allshire RC, Kouzarides T. 2001. Selective recognition of methylated lysine 9 on histone H3 by the HP1 chromo domain. Nature 410:120–124.

5. Schotta G, Ebert A, Krauss V, Fischer A, Hoffmann J, Rea S, Jenuwein T, Dorn R, Reuter G. 2002. Central role of Drosophila SU(VAR)3-9 in histone H3-K9 methylation and heterochromatic gene silencing. EMBO J 21:1121–1131.

6. Nakayama J, Rice JC, Strahl BD, Allis CD, Grewal SI. 2001. Role of histone H3 lysine 9 methylation in epigenetic control of heterochromatin assembly. Science 292:110–113.

7. Lomberk G, Wallrath L, Urrutia R. 2006. The Heterochromatin Protein 1 family. Genome Biol 7:228.

8. Kwon SH, Workman JL. 2008. The heterochromatin protein 1 (HP1) family: put away a bias toward HP1. Mol Cells 26:217–227.

9. Grewal SI, Jia S. 2007. Heterochromatin revisited. Nat Rev Genet 8:35–46.

10. Wang J, Jia ST, Jia S. 2016. New Insights into the Regulation of Heterochromatin. Trends Genet 32:284–294.

11. Zeng W, Ball AR, Jr., Yokomori K. 2010. HP1: heterochromatin binding proteins working the genome. Epigenetics 5:287–92.

12. Flueck C, Bartfai R, Volz J, Niederwieser I, Salcedo-Amaya AM, Alako BT, Ehlgen F, Ralph SA, Cowman AF, Bozdech Z, Stunnenberg HG, Voss TS. 2009. Plasmodium falciparum heterochromatin protein 1 marks genomic loci linked to phenotypic variation of exported virulence factors. PLoS Pathog 5:e1000569.

13. Perez-Toledo K, Rojas-Meza AP, Mancio-Silva L, Hernandez-Cuevas NA, Delgadillo DM, Vargas M, Martinez-Calvillo S, Scherf A, Hernandez-Rivas R. 2009. Plasmodium falciparum heterochromatin protein 1 binds to tri-methylated histone 3 lysine 9 and is linked to mutually exclusive expression of var genes. Nucleic Acids Res 37:2596–2606.

14. Brasher SV, Smith BO, Fogh RH, Nietlispach D, Thiru A, Nielsen PR, Broadhurst RW, Ball LJ, Murzina NV, Laue ED. 2000. The structure of mouse HP1 suggests a unique mode of single peptide recognition by the shadow chromo domain dimer. EMBO J 19:1587–1597.

15. Cowieson NP, Partridge JF, Allshire RC, McLaughlin PJ. 2000. Dimerisation of a chromo shadow domain and distinctions from the chromodomain as revealed by structural analysis. Curr Biol 10:517–525.

16. Lechner MS, Begg GE, Speicher DW, Rauscher FJ, III. 2000. Molecular determinants for targeting heterochromatin protein 1-mediated gene silencing: direct chromoshadow domain-KAP-1 corepressor interaction is essential. Mol Cell Biol 20:6449–6465.

17. Yamamoto K, Sonoda M. 2003. Self-interaction of heterochromatin protein 1 is required for direct binding to histone methyltransferase, SUV39H1. Biochem Biophys Res Commun 301:287–292.

18. Meehan RR, Kao CF, Pennings S. 2003. HP1 binding to native chromatin in vitro is determined by the hinge region and not by the chromodomain. EMBO J 22:3164–3174.

19. Muchardt C, Guilleme M, Seeler JS, Trouche D, Dejean A, Yaniv M. 2002. Coordinated methyl and RNA binding is required for heterochromatin localization of mammalian HP1alpha. EMBO Rep 3:975–981.

20. Mishima Y, Watanabe M, Kawakami T, Jayasinghe CD, Otani J, Kikugawa Y, Shirakawa M, Kimura H, Nishimura O, Aimoto S, Tajima S, Suetake I. 2013. Hinge and chromoshadow of HP1α participate in recognition of K9 methylated histone H3 in nucleosomes. J Mol Biol 425:54–70.

21. Nielsen AL, Oulad-Abdelghani M, Ortiz JA, Remboutsika E, Chambon P, Losson R. 2001. Heterochromatin formation in mammalian cells: interaction between histones and HP1 proteins. Mol Cell 7:729–739.

22. Horsley D, Hutchings A, Butcher GW, Singh PB. 1996. M32, a murine homologue of Drosophila heterochromatin protein 1 (HP1), localises to euchromatin within interphase nuclei and is largely excluded from constitutive heterochromatin. Cytogenet Cell Genet 73:308–311.

23. Eberhard L, Schneider S, Eiffler C, Kappel S, Giannakopoulos NN. 2015. Particle size distributions determined by optical scanning and by sieving in the assessment of masticatory performance of complete denture wearers. Clin Oral Investig 19:429–436.

24. Minc E, Courvalin JC, Buendia B. 2000. HP1gamma associates with euchromatin and heterochromatin in mammalian nuclei and chromosomes. Cytogenet Cell Genet 90:279–284.

25. Minc E, Allory Y, Worman HJ, Courvalin JC, Buendia B. 1999. Localization and phosphorylation of HP1 proteins during the cell cycle in mammalian cells. Chromosoma 108:220–234.

26. Wang G, Ma A, Chow CM, Horsley D, Brown NR, Cowell IG, Singh PB. 2000. Conservation of heterochromatin protein 1 function. Mol Cell Biol 20:6970–6983.

27. Powers JA, Eissenberg JC. 1993. Overlapping domains of the heterochromatin-associated protein HP1 mediate nuclear localization and heterochromatin binding. J Cell Biol 120:291–299.

28. Platero JS, Hartnett T, Eissenberg JC. 1995. Functional analysis of the chromo domain of HP1. EMBO J 14:3977–3986.

29. Witmer K, Fraschka SA, Vlachou D, Bartfai R, Christophides GK. 2020. An epigenetic map of malaria parasite development from host to vector. Sci Rep 10:6354.

30. Fraschka SA, Filarsky M, Hoo R, Niederwieser I, Yam XY, Brancucci NMB, Mohring F, Mushunje AT, Huang X, Christensen PR, Nosten F, Bozdech Z, Russell B, Moon RW, Marti M, Preiser PR, Bartfai R, Voss TS. 2018. Comparative Heterochromatin Profiling Reveals Conserved and Unique Epigenome Signatures Linked to Adaptation and Development of Malaria Parasites. Cell Host Microbe 23:407–420.

31. Allshire RC, Nimmo ER, Ekwall K, Javerzat JP, Cranston G. 1995. Mutations derepressing silent centromeric domains in fission yeast disrupt chromosome segregation. Genes Dev 9:218–233.

32. Nonaka N, Kitajima T, Yokobayashi S, Xiao G, Yamamoto M, Grewal SI, Watanabe Y. 2002. Recruitment of cohesin to heterochromatic regions by Swi6/HP1 in fission yeast. Nat Cell Biol 4:89–93.

33. Fischer T, Cui B, Dhakshnamoorthy J, Zhou M, Rubin C, Zofall M, Veenstra TD, Grewal SI. 2009. Diverse roles of HP1 proteins in heterochromatin assembly and functions in fission yeast. Proc Natl Acad Sci U S A 106:8998–9003.

34. Yi Q, Chen Q, Liang C, Yan H, Zhang Z, Xiang X, Zhang M, Qi F, Zhou L, Wang F. 2018. HP1 links centromeric heterochromatin to centromere cohesion in mammals. EMBO Rep 19.

35. Salcedo-Amaya AM, van Driel MA, Alako BT, Trelle MB, van den Elzen AM, Cohen AM, Janssen-Megens EM, van D, V, Selzer RR, Iniguez AL, Green RD, Sauerwein RW, Jensen ON, Stunnenberg HG. 2009. Dynamic histone H3 epigenome marking during the intraerythrocytic cycle of Plasmodium falciparum. Proc Natl Acad Sci U S A 106:9655–9660.

36. Lopez-Rubio JJ, Mancio-Silva L, Scherf A. 2009. Genome-wide analysis of heterochromatin associates clonally variant gene regulation with perinuclear repressive centers in malaria parasites. Cell Host Microbe 5:179–190.

37. Baruch DI, Pasloske BL, Singh HB, Bi X, Ma XC, Feldman M, Taraschi TF, Howard RJ. 1995. Cloning the P. falciparum gene encoding PfEMP1, a malarial variant antigen and adherence receptor on the surface of parasitized human erythrocytes. Cell 82:77–87.

38. Su XZ, Heatwole VM, Wertheimer SP, Guinet F, Herrfeldt JA, Peterson DS, Ravetch JA, Wellems TE. 1995. The large diverse gene family var encodes proteins involved in cytoadherence and antigenic variation of Plasmodium falciparum-infected erythrocytes. Cell 82:89–100.

39. Smith JD, Chitnis CE, Craig AG, Roberts DJ, Hudson-Taylor DE, Peterson DS, Pinches R, Newbold CI, Miller LH. 1995. Switches in expression of Plasmodium falciparum var genes correlate with changes in antigenic and cytoadherent phenotypes of infected erythrocytes. Cell 82:101–110.

40. Gardner MJ, Hall N, Fung E, White O, Berriman M, Hyman RW, Carlton JM, Pain A, Nelson KE, Bowman S, Paulsen IT, James K, Eisen JA, Rutherford K, Salzberg SL, Craig A, Kyes S, Chan MS, Nene V, Shallom SJ, Suh B, Peterson J, Angiuoli S, Pertea M, Allen J, Selengut J, Haft D, Mather MW, Vaidya AB, Martin DM, Fairlamb AH, Fraunholz MJ, Roos DS, Ralph SA, McFadden GI, Cummings LM, Subramanian GM, Mungall C, Venter JC, Carucci DJ, Hoffman SL, Newbold C, Davis RW, Fraser CM, Barrell B. 2002. Genome sequence of the human malaria parasite Plasmodium falciparum. Nature 419:498–511.

41. Scherf A, Lopez-Rubio JJ, Riviere L. 2008. Antigenic variation in Plasmodium falciparum. Annu Rev Microbiol 62:445–470.

42. Hviid L, Jensen AT. 2015. PfEMP1, A Parasite Protein Family of Key Importance in Plasmodium falciparum Malaria Immunity and Pathogenesis. Adv Parasitol 88:51–84.

43. Wahlgren M, Goel S, Akhouri RR. 2017. Variant surface antigens of Plasmodium falciparum and their roles in severe malaria. Nat Rev Microbiol 15:479–491.

44. Scherf A, Hernandez-Rivas R, Buffet P, Bottius E, Benatar C, Pouvelle B, Gysin J, Lanzer M. 1998. Antigenic variation in malaria: in situ switching, relaxed and mutually exclusive transcription of var genes during intra-erythrocytic development in Plasmodium falciparum. EMBO J 17:5418–5426.

45. Guizetti J, Scherf A. 2013. Silence, activate, poise and switch! Mechanisms of antigenic variation in Plasmodium falciparum. Cell Microbiol 15:718–726.

46. Deitsch KW, Dzikowski R. 2017. Variant Gene Expression and Antigenic Variation by Malaria Parasites. Annu Rev Microbiol 71:625–641.

47. Lopez-Rubio JJ, Gontijo AM, Nunes MC, Issar N, Hernandez RR, Scherf A. 2007. 5’ flanking region of var genes nucleate histone modification patterns linked to phenotypic inheritance of virulence traits in malaria parasites. Mol Microbiol 66:1296–1305.

48. Chookajorn T, Dzikowski R, Frank M, Li F, Jiwani AZ, Hartl DL, Deitsch KW. 2007. Epigenetic memory at malaria virulence genes. Proc Natl Acad Sci U S A 104:899–902.

49. Brancucci NMB, Bertschi NL, Zhu L, Niederwieser I, Chin WH, Wampfler R, Freymond C, Rottmann M, Felger I, Bozdech Z, Voss TS. 2014. Heterochromatin protein 1 secures survival and transmission of malaria parasites. Cell Host Microbe 16:165–176.

50. Lopez-Rubio JJ, Mancio-Silva L, Scherf A. 2009. Genome-wide analysis of heterochromatin associates clonally variant gene regulation with perinuclear repressive centers in malaria parasites. Cell Host Microbe 5:179–90.

51. Freitas-Junior LH, Hernandez-Rivas R, Ralph SA, Montiel-Condado D, Ruvalcaba-Salazar OK, Rojas-Meza AP, Mancio-Silva L, Leal-Silvestre RJ, Gontijo AM, Shorte S, Scherf A. 2005. Telomeric heterochromatin propagation and histone acetylation control mutually exclusive expression of antigenic variation genes in malaria parasites. Cell 121:25–36.

52. Duraisingh MT, Voss TS, Marty AJ, Duffy MF, Good RT, Thompson JK, Freitas-Junior LH, Scherf A, Crabb BS, Cowman AF. 2005. Heterochromatin silencing and locus repositioning linked to regulation of virulence genes in Plasmodium falciparum. Cell 121:13–24.

53. Ralph SA, Scheidig-Benatar C, Scherf A. 2005. Antigenic variation in Plasmodium falciparum is associated with movement of var loci between subnuclear locations. Proc Natl Acad Sci U S A 102:5414–5419.

54. Voss TS, Healer J, Marty AJ, Duffy MF, Thompson JK, Beeson JG, Reeder JC, Crabb BS, Cowman AF. 2006. A var gene promoter controls allelic exclusion of virulence genes in Plasmodium falciparum malaria. Nature 439:1004–1008.

55. Volz JC, Bartfai R, Petter M, Langer C, Josling GA, Tsuboi T, Schwach F, Baum J, Rayner JC, Stunnenberg HG, Duffy MF, Cowman AF. 2012. PfSET10, a Plasmodium falciparum methyltransferase, maintains the active var gene in a poised state during parasite division. Cell Host Microbe 11:7–18.

56. Dzikowski R, Li F, Amulic B, Eisberg A, Frank M, Patel S, Wellems TE, Deitsch KW. 2007. Mechanisms underlying mutually exclusive expression of virulence genes by malaria parasites. EMBO Rep 8:959–965.

57. Kafsack BF, Rovira-Graells N, Clark TG, Bancells C, Crowley VM, Campino SG, Williams AE, Drought LG, Kwiatkowski DP, Baker DA, Cortes A, Llinas M. 2014. A transcriptional switch underlies commitment to sexual development in malaria parasites. Nature 507:248–252.

58. Sinha A, Hughes KR, Modrzynska KK, Otto TD, Pfander C, Dickens NJ, Religa AA, Bushell E, Graham AL, Cameron R, Kafsack BF, Williams AE, Llinas M, Berriman M, Billker O, Waters AP. 2014. A cascade of DNA-binding proteins for sexual commitment and development in Plasmodium. Nature 507:253–257.

59. Boltryk SD, Passecker A, Alder A, van de Vegte-Bolmer M, Sauerwein RW, Brancucci NMB, Beck H-P, Gilberger T-W, Voss TS. 2020. CRISPR/Cas9-engineered inducible gametocyte producer lines as a novel tool for basic and applied research on Plasmodium falciparum> malaria transmission stages. bioRxiv doi:10.1101/2020.10.05.326868:2020.10.05.326868.

60. Balaji S, Babu MM, Iyer LM, Aravind L. 2005. Discovery of the principal specific transcription factors of Apicomplexa and their implication for the evolution of the AP2-integrase DNA binding domains. Nucleic Acids Res 33:3994–4006.

61. Filarsky M, Fraschka SA, Niederwieser I, Brancucci NMB, Carrington E, Carrio E, Moes S, Jenoe P, Bartfai R, Voss TS. 2018. GDV1 induces sexual commitment of malaria parasites by antagonizing HP1-dependent gene silencing. Science 359:1259–1263.

62. Josling GA, Russell TJ, Venezia J, Orchard L, van Biljon R, Painter HJ, Llinas M. 2020. Dissecting the role of PfAP2-G in malaria gametocytogenesis. Nat Commun 11:1503.

63. Llora-Batlle O, Michel-Todo L, Witmer K, Toda H, Fernandez-Becerra C, Baum J, Cortes A. 2020. Conditional expression of PfAP2-G for controlled massive sexual conversion in Plasmodium falciparum. Sci Adv 6:eaaz5057.

64. Bancells C, Llora-Batlle O, Poran A, Notzel C, Rovira-Graells N, Elemento O, Kafsack BFC, Cortes A. 2019. Revisiting the initial steps of sexual development in the malaria parasite Plasmodium falciparum. Nat Microbiol 4:144–154.

65. Kent RS, Modrzynska KK, Cameron R, Philip N, Billker O, Waters AP. 2018. Inducible developmental reprogramming redefines commitment to sexual development in the malaria parasite Plasmodium berghei. Nat Microbiol 3:1206–1213.

66. Eksi S, Morahan BJ, Haile Y, Furuya T, Jiang H, Ali O, Xu H, Kiattibutr K, Suri A, Czesny B, Adeyemo A, Myers TG, Sattabongkot J, Su XZ, Williamson KC. 2012. Plasmodium falciparum gametocyte development 1 (Pfgdv1) and gametocytogenesis early gene identification and commitment to sexual development. PLoS Pathog 8:e1002964.

67. Brancucci NMB, Gerdt JP, Wang C, De Niz M, Philip N, Adapa SR, Zhang M, Hitz E, Niederwieser I, Boltryk SD, Laffitte MC, Clark MA, Gruring C, Ravel D, Blancke Soares A, Demas A, Bopp S, Rubio-Ruiz B, Conejo-Garcia A, Wirth DF, Gendaszewska-Darmach E, Duraisingh MT, Adams JH, Voss TS, Waters AP, Jiang RHY, Clardy J, Marti M. 2017. Lysophosphatidylcholine Regulates Sexual Stage Differentiation in the Human Malaria Parasite Plasmodium falciparum. Cell 171:1532–1544 e15.

68. Jones ML, Das S, Belda H, Collins CR, Blackman MJ, Treeck M. 2016. A versatile strategy for rapid conditional genome engineering using loxP sites in a small synthetic intron in Plasmodium falciparum. Sci Rep 6:21800.

69. Collins CR, Das S, Wong EH, Andenmatten N, Stallmach R, Hackett F, Herman JP, Muller S, Meissner M, Blackman MJ. 2013. Robust inducible Cre recombinase activity in the human malaria parasite Plasmodium falciparum enables efficient gene deletion within a single asexual erythrocytic growth cycle. Mol Microbiol 88:687–701.

70. Bui HTN, Niederwieser I, Bird MJ, Dai W, Brancucci NMB, Moes S, Jenoe P, Lucet IS, Doerig C, Voss TS. 2019. Mapping and functional analysis of heterochromatin protein 1 phosphorylation in the malaria parasite Plasmodium falciparum. Sci Rep 9:16720.

71. Flueck C, Bartfai R, Niederwieser I, Witmer K, Alako BT, Moes S, Bozdech Z, Jenoe P, Stunnenberg HG, Voss TS. 2010. A major role for the Plasmodium falciparum ApiAP2 protein PfSIP2 in chromosome end biology. PLoS Pathog 6:e1000784.

72. Brameier M, Krings A, MacCallum RM. 2007. NucPred--predicting nuclear localization of proteins. Bioinformatics 23:1159–60.

73. Ponnudurai T, Lensen AH, Meis JF, Meuwissen JH. 1986. Synchronization of Plasmodium falciparum gametocytes using an automated suspension culture system. Parasitology 93 (Pt 2):263–274.

74. Fivelman QL, McRobert L, Sharp S, Taylor CJ, Saeed M, Swales CA, Sutherland CJ, Baker DA. 2007. Improved synchronous production of Plasmodium falciparum gametocytes in vitro. Mol Biochem Parasitol 154:119–123.

75. Bruce MC, Carter RN, Nakamura K, Aikawa M, Carter R. 1994. Cellular location and temporal expression of the Plasmodium falciparum sexual stage antigen Pfs16. Mol Biochem Parasitol 65:11–22.

76. Yamada T, Fukuda R, Himeno M, Sugimoto K. 1999. Functional domain structure of human heterochromatin protein HP1(Hsalpha): involvement of internal DNA-binding and C-terminal self-association domains in the formation of discrete dots in interphase nuclei. J Biochem 125:832–837.

77. Smothers JF, Henikoff S. 2001. The hinge and chromo shadow domain impart distinct targeting of HP1-like proteins. Mol Cell Biol 21:2555–2569.

78. Oehring SC, Woodcroft BJ, Moes S, Wetzel J, Dietz O, Pulfer A, Dekiwadia C, Maeser P, Flueck C, Witmer K, Brancucci NM, Niederwieser I, Jenoe P, Ralph SA, Voss TS. 2012. Organellar proteomics reveals hundreds of novel nuclear proteins in the malaria parasite Plasmodium falciparum. Genome Biol 13:R108.

79. Cui L, Fan Q, Cui L, Miao J. 2008. Histone lysine methyltransferases and demethylases in Plasmodium falciparum. Int J Parasitol 38:1083–1097.

80. Volz J, Carvalho TG, Ralph SA, Gilson P, Thompson J, Tonkin CJ, Langer C, Crabb BS, Cowman AF. 2010. Potential epigenetic regulatory proteins localise to distinct nuclear sub-compartments in Plasmodium falciparum. Int J Parasitol 40:109–121.

81. Tibúrcio M, Hitz E, Niederwieser I, Kelly G, Davies H, Doerig C, Billker O, Voss TS, Treeck M. 2020. GDV1 C-terminal truncation of 39 amino acids disrupts sexual commitment in Plasmodium falciparum. bioRxiv doi:10.1101/2020.10.28.360123:2020.10.28.360123.

82. Broadbent KM, Broadbent JC, Ribacke U, Wirth D, Rinn JL, Sabeti PC. 2015. Strand-specific RNA sequencing in Plasmodium falciparum malaria identifies developmentally regulated long non-coding RNA and circular RNA. BMC Genomics 16:454.

83. Lambros C, Vanderberg JP. 1979. Synchronization of Plasmodium falciparum erythrocytic stages in culture. J Parasitol 65:418–420.

84. Knuepfer E, Napiorkowska M, van Ooij C, Holder AA. 2017. Generating conditional gene knockouts in Plasmodium - a toolkit to produce stable DiCre recombinase-expressing parasite lines using CRISPR/Cas9. Sci Rep 7:3881.

85. Witmer K, Schmid CD, Brancucci NM, Luah YH, Preiser PR, Bozdech Z, Voss TS. 2012. Analysis of subtelomeric virulence gene families in Plasmodium falciparum by comparative transcriptional profiling. Mol Microbiol 84:243–259.

