## Supplementary Information for "Investigation of heterochromatin protein 1 function in the malaria parasite *Plasmodium falciparum* using a conditional domain deletion and swapping approach"

This PDF file includes:

Supplementary Methods

Supplementary Figures S1 and S2

Supplementary Table S1

### Supplementary Methods

#### Generation of the pBcam-ΔHinge-3HA-Cherry, pBcam-hyb-PbHinge-3HA-Cherry and pBcam-hyb-PbCSD-3HA-Cherry plasmids

The pBcam-ΔHinge-3HA-Cherry plasmid was constructed as follows. The fragment PfCD spanning bps +1 to +222 of the recodonised *pfhp1* sequence was amplified from a pUC57 plasmid containing a synthetic recodonized *pfhp1* coding sequence (pUC57-re-*pfhp1)* (1) using primers F11 and R148. The fragment PfSIP2.linker (spanning bps +694 to +762 of the *pfsip2* coding sequence) was amplified from 3D7 gDNA using primers F4 and R5. The fragment PfCSD spanning bps +532 to +798 of the recodonized *pfhp1* sequence and omitting the stop codon was amplified from pUC57-re-*pfhp1* using primers F2 and R3. A hybrid fragment consisting of the PfSIP2.linker and PfCSD fragments was amplified by fusion PCR from a mixture of the PfSIP2.linker and PfCSD PCR templates using primers F4 and R3. The final hybrid fragment of PfCD/PfSIP2.linker/PfCSD was amplified by fusion PCR from a mixture of PfCD and PfSIP2.linker/PfCSD PCR templates using primers F11 and R3 and cloned into pBcam-3HA-Cherry (2) after digestion with *BamH*I and *Nhe*I and ligation with T4 DNA ligase.

The pBcam-hyb-PbHinge-3HA-Cherry plasmid was constructed as follows. The fragment PfCD spanning bps +1 to +222 of the recodonized *pfhp1* sequence was amplified from pUC57-re-*pfhp1* using primers F11 and R1. The fragment PbHinge spanning bps +223 to +576 of the *pbhp1* sequence was amplified from *P. berghei* gDNA using primers F35 and R36. The fragment PfCSD spanning bps +532 to +798 of the recodonized *pfhp1* sequence and omitting stop codon was amplified from pUC57-re-*pfhp1* using primers F2 and R42. A hybrid fragment consisting of the PfCD/PbHinge fragments was amplified by fusion PCR from a mixture of PfCD and PbHinge PCR templates using primers F11 and R36. The final hybrid fragment of PfCD/PbHinge/PfCSD was amplified by fusion PCR from a mixture of the PfCD/PbHinge and PfCSD PCR templates using primers F11 and R42 and cloned into pBcam-3HA-Cherry after digestion with *BamH*I and *Not*I and ligation with T4 DNA ligase.

The pBcam-hyb-PbCSD-3HA-Cherry plasmid was constructed as follows. The fragment PfCD.Hinge spanning bps +1 to +570 of the recodonized *pfhp1* sequence was amplified from pUC57-re-*pfhp1* using primers F11 and R12. The fragment PbCSD spanning bps +616 to +843 of the *pbhp1* sequence and omitting stop codon was amplified from *P. berghei* gDNA using primers F38 and R41. A hybrid fragment consisting of PfCD.Hinge/PbCSD was amplified by fusion PCR from a mixture of the PfCD.Hinge and PbCSD PCR templates using primers F11 and R41 and cloned into pBcam-3HA-Cherry after digestion with *BamH*I and *Not*I and ligation with T4 DNA ligase.

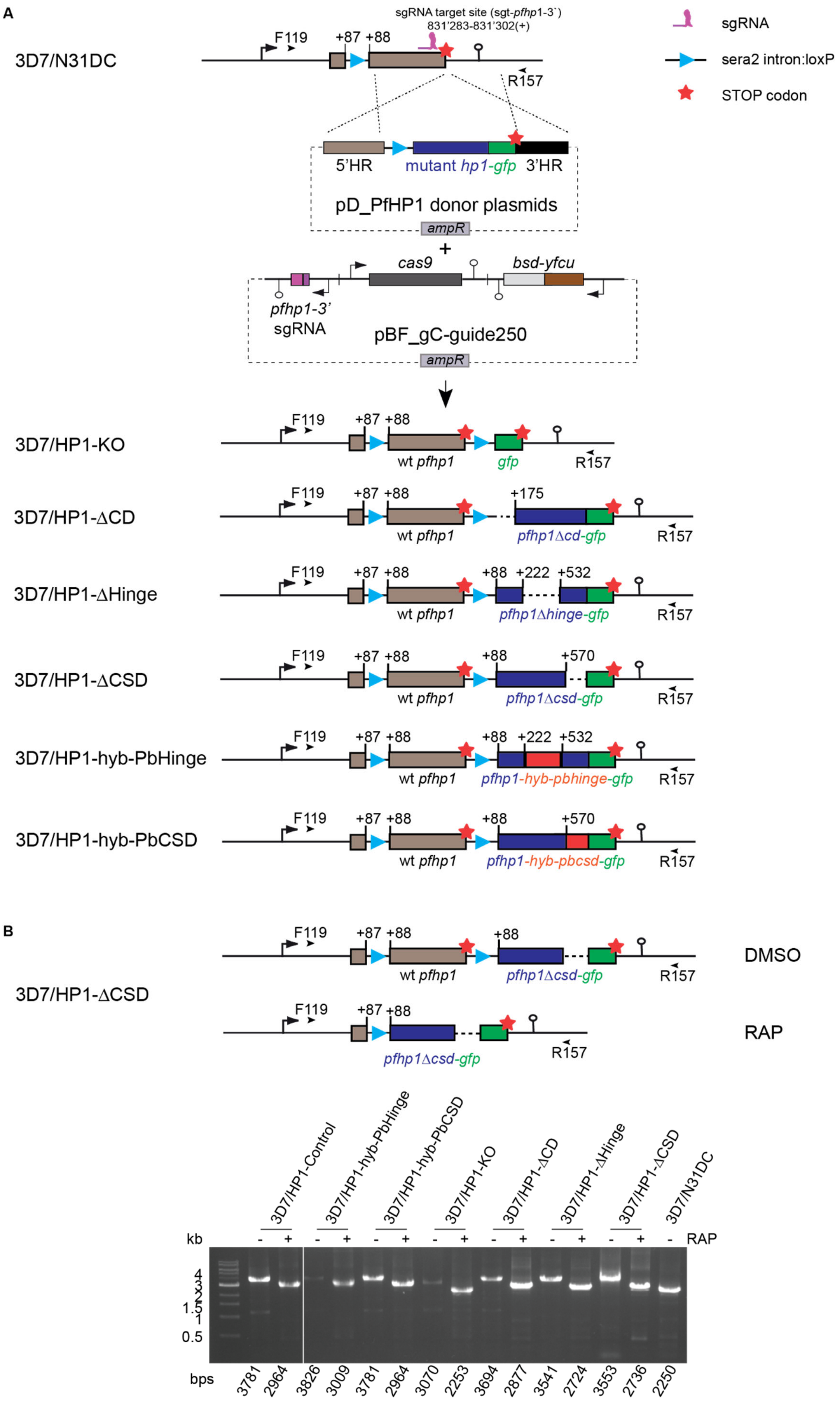

A
sgRNA target site (sgt-pfhp1-3`)
831'283-831'302(+)
sgRNA
sera2 intron:loxP
STOP codon
3D7/N31DC
F119
+87 +88
R157
5'HR
mutant hp1-gfp
3'HR
pD_PfHP1 donor plasmids
ampR
+
cas9
bsd-yfcu
pfhp1-3'
sgRNA
pBF_gC-guide250
ampR
3D7/HP1-KO
wt pfhp1
gfp
3D7/HP1-ΔCD
+175
pfhp1Δcd-gfp
3D7/HP1-ΔHinge
+88 +222 +532
pfhp1Δhinge-gfp
3D7/HP1-ΔCSD
+88 +570
pfhp1Δcsd-gfp
3D7/HP1-hyb-PbHinge
pfhp1-hyb-pbhinge-gfp
3D7/HP1-hyb-PbCSD
pfhp1-hyb-pbcsd-gfp
B
3D7/HP1-ΔCSD
DMSO
RAP
3D7/HP1-Control
3D7/HP1-hyb-PbHinge
3D7/HP1-hyb-PbCSD
3D7/HP1-KO
3D7/HP1-ΔCD
3D7/HP1-ΔHinge
3D7/HP1-ΔCSD
3D7/N31DC
kb
RAP
4
3
2
1.5
1
0.5
bps
3781 2964 3826 3009 3781 2964 3070 2253 3694 2877 3541 2724 3553 2736 2250

**Supplementary Figure 1. CRISPR/Cas9-based gene editing strategy to generate DiCre-inducible PfHP1 truncation and PfHP1-PbHP1 hybrid mutants. (A)** Top: schematic map of the *pfhp1* locus (PF3D7_1220900) in 3D7/N31DC parasites. 3D7/N31DC parasites have previously been obtained by inserting a 103 bp *sera2* intron:loxP element (3) (light blue triangle) into the 5' end of the *pfhp1* gene in 3D7/1G5DiCre parasites (4) using CRISPR/Cas9-based gene editing (1). The nucleotide positions of the sgt_*pfhp1*-3' sgRNA target sequence is indicated (chromosome 12 coordinates). Center: Schematic maps of a pD_PfHP1 donor plasmid co-transfected with the pBF_gC-guide250 CRISPR/Cas9 transfection vector (1). The pD_PfHP1 donor plasmids contain an assembly of the 103 bp *sera2* intron:loxP element (light blue triangle) and a mutated *pfhp1* sequence (dark blue) fused to *gfp* (green) flanked by 5' and 3' homology regions (HR) (sandstone, black) for homology-directed repair. The pBF_gC-guide250 plasmid contains expression cassettes for SpCas9 (dark grey), the sgRNA (purple) and the *bsd-yfcu* fusion selection marker (light grey-brown). Bottom: Schematic maps of the modified *pfhp1* loci after CRISPR/Cas9-based gene editing in 3D7/HP1-KO, 3D7/HP1-ΔCD, 3D7/HP1-ΔHinge, 3D7/HP1-ΔCSD, 3D7/HP1-hyb-PbHinge and 3D7/HP1-hyb-PbCSD are shown. Light blue triangles represent *sera2* intron:loxP elements, red stars represent STOP codons. Brown and blue boxes represent the wild type and the replacing mutant *pfhp1* sequences, respectively. Numbers refer to the nucleotide position within the *pfhp1* coding sequence. The black arrowheads indicate the binding sites of the F119 and R157 PCR primers used to confirm correct gene editing of the *pfhp1* locus and efficient DiCre-mediated excision upon RAP treatment. **(B)** PCR confirmation of correct editing of the *pfhp1* locus and efficient excision of the floxed *pfhp1* gene after RAP treatment. The schematic on top shows the *pfhp1* locus before (DMSO) and after RAP treatment (RAP) in the 3D7/HP1-ΔCSD parasite line as an example. PCRs were performed on gDNA of DMSO-treated (-) and RAP-treated (+) 3D7/HP1-Control (1), 3D7/HP1-hyb-PbHinge, 3D7/HP1-hyb-PbCSD, 3D7/HP1-KO, 3D7/HP1-ΔCD, 3D7/HP1-ΔHinge, 3D7/HP1-ΔCSD cell lines using the primer combination F119 and R157. The length of PCR fragments amplified from the correctly edited *pfhp1* locus before (-) and after excision upon RAP treatment (+) from the various PfHP1 mutant parasite lines and the 3D7/N31DC mother line are indicated at the bottom of the figure. The correct excision of floxed DNA after RAP treatment (+) results in a decreased fragment size compared to the DMSO-treated (-) control condition.

```
                                  Chromo domain
PfHP1        MTGSDEEFEIGDILEIKKKKNGFIYLVKWKGYSDDENTWEPESNLIHLTTFKKKMESLKT      60
PbHP1        MTGSDEEFEIGDILDVKRKKNGFIYLVKWKGYSDDENTWEPESNLLHLTDFKKKMEYLKS      60
             **************::*:***************************:*** ****** **:
                    #                    #  #     #   # #

PfHP1        NFLSKANETNGDGKILKNHILAPTQED----DSIKSKGRSSLAPRRKMSRKSLTNKLEN-      115
PbHP1        IYLNKIDRTSSDSKIMKKNNVQLFDQDDMGNTLMKPKGRTTLISRKRGHKRGMRNRMRNR      120
              :*.* :.*..*.**:*:: :   ::*      :* ***::*  *::  ::.: *::.*

PfHP1        --------KKNLSLSDNSLKKSDEEDNESVKHENHVN--DGNLLNVEDVYSVRIKNKKLE      165
PbHP1        MRNRIGNKSSASSVTDGSLKKSDDDDNQSIKKESSSNNYNNTLLNIEDVYSVRIKNRKME      180
                     ..  *::*.******::**:*:*:*.  *  :..***:**********:*:*
                                     Chromoshadow domain
PfHP1        FLASLKNESPQWVEETNIRRTGHLNIKVNDFKRYVRRKKSSRGNRIVIKNLHNVGDELYI      225
PbHP1        FLASLKNASPQWVEESNIRSTGHLNIKVNDFKKYIKRKKTSKGSRIVIKNLHNVGDELYI      240
             ******* *******:*** ************:*::***:*:*.****************

PfHP1        SVIHNINNKEIHSLYPSKVIEYIYPQELLNFLLSRLRYRTA      266
PbHP1        SVIHNINNKEIHSLYPSKVIEYIYPQELLNFLLSRLRYRTV      281
             ****************************************.
                   #        #   #   #    # #
                                 ##   ## #
```

**Supplementary Figure S2. Pairwise sequence alignment of PfHP1 and PbHP1**. Alignment of the PfHP1 and PbHP1 amino acid sequences by EMBOSS Needle (https://www.ebi.ac.uk/Tools/psa/emboss_needle/). Identical amino acids are marked by asterisks. The PfHP1 CD and CSD are highlighted by horizontal lines; purple hashtags denote residues important for H3K9me binding by the CD; blue and green hashtags denote residues important for CSD homo- and hetero-dimerisation, respectively (5).

**Supplementary Table S1. Oligonucleotides used in this study.**

| Application | Oligo name | Sequence (5'-3') |
|---|---|---|
| PCR for the cloning of transfection vectors | **R1** | tttaccatctccatttgtttcatttg |
| | **F2** | gttgaagaaacaaatattagaagaac |
| | **R3** | cagtgctagctgctgttctatatcttaatcttg |
| | **F4** | gaaacaaatggagatggtaaaggaataacctttgaacaattatatc |
| | **R5** | ctaatatttgtttcttcaacacctacattttcaaatactcgtac |
| | **F11** | cagtggatccaaaaaatgacaggtagtgatgaag |
| | **R12** | attcaaatgacctgttcttc |
| | **F35** | caaatgaaacaaatggagatggtaaaattatgaaaaaaaacaatgtacag |
| | **R36** | gttcttctaatatttgtttcttcaacccattgtggtgatgcattttttaag |
| | **F38** | gaagaacaggtcatttgaatataaaagttaacgattttaaaaaa |
| | **R41** | caatgcggccgcaaccgttctatatctaagtcttg |
| | **R42** | caatgcggccgctgctgttctatatcttaatcttg |
| | **F139** | gtaaataaaaaaaataatatacaataac |
| | **R143** | ctaaaagaatataaaatatataaatat |
| | **R148** | agggtatcaccttcaaacttgacttcagcacgtgtcttgtag |
| | **F158** | cgttggccgattcattaatgaaaggatattcagatgatgag |
| | **R159** | gttattgtatattattttttttatttacttacgctgttctatatcttaatc |
| | **F160** | atatttatatattttatattcttttagaaaggctattcagatgatg |
| | **R161** | gttcttctcctttactcataaccgttctatatctaagtc |
| | **F162** | atgagtaaaggagaagaac |
| | **R163** | cctcttcgctattacgccaggaggttaaaattctaaactatatg |
| | **F164** | atatttatatattttatattcttttagaaaggatatagtgatgatga |
| | **R165** | gttcttctcctttactcattgctgttctatatcttaatc |
| | **F177** | atatttatatattttatattcttttagaaaacaaatttcttatctaaag |
| | **R178** | gttcttctcctttactcatattcaaatgacctgttcttc |
| | **PCRA_F** | ctggcgtaatagcgaagagg |
| | **PCRA_R** | cattaatgaatcggccaacg |
| Diagnostic PCR on gDNA | **F119** | gtgtgtgtttaagaaaaaatatg |
| | **R157** | catgtagccaaaatatgtg |

### References


1. Bui HTN, Niederwieser I, Bird MJ, Dai W, Brancucci NMB, Moes S, Jenoe P, Lucet IS, Doerig C, Voss TS. 2019. Mapping and functional analysis of heterochromatin protein 1 phosphorylation in the malaria parasite Plasmodium falciparum. Sci Rep 9:16720.

2. Witmer K, Schmid CD, Brancucci NM, Luah YH, Preiser PR, Bozdech Z, Voss TS. 2012. Analysis of subtelomeric virulence gene families in Plasmodium falciparum by comparative transcriptional profiling. Mol Microbiol 84:243-259.

3. Jones ML, Das S, Belda H, Collins CR, Blackman MJ, Treeck M. 2016. A versatile strategy for rapid conditional genome engineering using loxP sites in a small synthetic intron in Plasmodium falciparum. Sci Rep 6:21800.

4. Collins CR, Das S, Wong EH, Andenmatten N, Stallmach R, Hackett F, Herman JP, Muller S, Meissner M, Blackman MJ. 2013. Robust inducible Cre recombinase activity in the human malaria parasite Plasmodium falciparum enables efficient gene deletion within a single asexual erythrocytic growth cycle. Mol Microbiol 88:687-701.

5. Flueck C, Bartfai R, Volz J, Niederwieser I, Salcedo-Amaya AM, Alako BT, Ehlgen F, Ralph SA, Cowman AF, Bozdech Z, Stunnenberg HG, Voss TS. 2009. Plasmodium falciparum heterochromatin protein 1 marks genomic loci linked to phenotypic variation of exported virulence factors. PLoS Pathog 5:e1000569.
